# *Lotus japonicus* karrikin receptors display divergent ligand-binding specificities and organ-dependent redundancy

**DOI:** 10.1101/754937

**Authors:** Samy Carbonnel, Salar Torabi, Maximilian Griesmann, Elias Bleek, Yuhong Tang, Stefan Buchka, Veronica Basso, Mitsuru Shindo, François-Didier Boyer, Trevor L. Wang, Michael Udvardi, Mark Waters, Caroline Gutjahr

**Affiliations:** LMU Munich, Faculty of Biology, Genetics, Biocenter Martinsried, Großhaderner Str. 2-4, 82152 Martinsried, Germany; Technical University of Munich (TUM), School of Life Sciences Weihenstephan, Plant Genetics, Emil Ramann Str. 4, 85354 Freising, Germany; Noble Research Institute, Ardmore, Oklahoma 73401 USA; Institute for Materials Chemistry and Engineering, Kyushu University, Kasugakoen-6, Kasuga, Fukuoka, 816-8580, Japan; Université Paris-Saclay, CNRS, Institut de Chimie des Substances Naturelles, UPR 2301, 91198, Gif-sur-Yvette, France; John Innes Centre, Norwich Research Park, Norwich NR4 7UH, UK; School of Molecular Sciences, The University of Western Australia, 35 Stirling Hwy, Perth, WA 6009, Australia; Australian Research Council Centre of Excellence in Plant Energy Biology, The University of Western Australia, 35 Stirling Hwy, Perth, WA 6009, Australia

**Author notes:** Department of Biology, University of Fribourg, Chemin du Musée 10, 1700 Fribourg, Switzerland (Samy Carbonnel). These authors contributed equally to this work (Samy Carbonnel, Salar Torabi).

## Abstract

Karrikins (KARs), smoke-derived butenolides, are perceived by the α/β-fold hydrolase KARRIKIN INSENSITIVE2 (KAI2) and are thought to mimic endogenous, yet elusive plant hormones tentatively called KAI2-ligands (KLs). The sensitivity to different karrikin types as well as the number of *KAI2* paralogs varies among plant species, suggesting diversification and co-evolution of ligand-receptor relationships. In legumes, which comprise a number of important crops with protein-rich, nutritious seed, *KAI2* has duplicated. We report sub-functionalization of KAI2a and KAI2b in the model legume *Lotus japonicus* and demonstrate that their ability to bind the synthetic ligand GR24^*ent*-5DS^ differs *in vitro* as well as in genetic assays in *Lotus japonicus* and in the heterologous *Arabidopsis thaliana* background. These differences can be explained by the exchange of a widely conserved phenylalanine in the binding pocket of KAI2a with a tryptophan in KAI2b, which occured independently in KAI2 proteins of several unrelated angiosperms. Furthermore, two polymorphic residues in the binding pocket are conserved across a number of legumes and may contribute to ligand binding preferences. Unexpectedly, *L. japonicus* responds to diverse synthetic KAI2-ligands in an organ-specific manner. Hypocotyl development responds to KAR_1_, KAR_2_ and *rac*-GR24, while root system development responds only to KAR_1_. This organ-specificity cannot be explained by receptor-ligand preferences alone, because *Lj*KAI2a is sufficient for karrikin responses in the hypocotyl, while *Lj*KAI2a and *Lj*KAI2b operate redundantly in roots. Our findings open novel research avenues into the evolution and diversity of butenolide ligand-receptor relationships, their ecological significance and the mechanisms controlling diverse developmental responses to different KAI2 ligands.

## Introduction

Karrikins (KARs) are small butenolide compounds derived from smoke of burning vegetation that were identified as germination stimulants of fire-following plants [1]. They can also accelerate seed germination of species that do not grow in fire-prone environments such as *Arabidopsis thaliana*, which enabled the identification of genes encoding karrikin receptor components via forward and reverse genetics. The α/β-fold hydrolase KARRIKIN INSENSITIVE2 (KAI2) is thought to bind KARs, and interacts with the F-box protein MORE AXILLIARY GROWTH 2 (MAX2) that is required for ubiquitylation of repressor proteins via the Skp1-Cullin-F-box (SCF) complex [2–9]. There are six known KARs, of which KAR_1_ is most abundant in smoke-water and most active on seed germination of fire-following plants [1, 10, 11], but Arabidopsis responds more strongly to KAR_2_, which lacks the methyl group at the butenolide ring that is characteristic for KAR_1_ [2, 3, 11]. Both KAR_1_ and KAR_2_ are commercially available and commonly used in research.

Arabidopsis KAI2 regulates several traits in addition to seed germination, including light-dependent hypocotyl growth inhibitition, cotyledon and rosette leaf area, cuticle thickness, root hair length and density, root skewing and lateral root density [4, 12–15]. Moreover, the rice orthologs of *KAI2* (*D14-LIKE*) and *MAX2* (*D3*) are essential for root colonization by arbuscular mycorrhiza (AM) fungi, and are involved in regulating mesocotyl elongation [7, 16, 17]. These roles of KAI2, unrelated to smoke and seed germination, suggest that karrikins mimic yet-unknown endogenous (and possibly AM fungus-derived) signalling molecules that bind to KAI2 to regulate plant development or AM symbiosis, and are provisionally called KAI2-ligands (KLs) [12, 18].

Structurally, KARs resemble the apocarotenoid strigolactones (SLs), which were originally discovered in root exudates in the rhizosphere [19], where they act as germination cues for parasitic weeds [19] and as stimulants of AM fungi [20, 21]. In addition to their function in the rhizosphere, SLs function endogenously as phytohormones and repress shoot branching [22, 23]. SL signalling also affects secondary growth; and co-regulates lateral and adventitious root formation and rice mesocotyl elongation with the karrikin signalling pathway [7, 15, 24, 25]. As with KARs, SLs are perceived by an α/β-fold hydrolase D14/DAD2 that, like KAI2, depends for function on a serine–histidine–aspartate catalytic triad within the ligand binding pocket [26, 27]. As KAI2, D14 interacts with the SCF-complex via the same F-box protein MAX2 [9, 26] to ubiquitylate repressors of the SMXL family and mark them for degradation by the 26S proteasome [28–31].

Phylogenetic analysis of the α/β-fold hydrolase receptors in extant land plants revealed that an ancestral *KAI2* is already present in charophyte algae, while the so-called eu-*KAI2* is ubiquitous among land plants. The strigolactone receptor gene, *D14* evolved only in the seed plants likely through duplication of *KAI2* and sub-functionalization [32]. An additional duplication in the seed plants gave rise to *D14-LIKE2* (*DLK2*), an α/β-fold hydrolase of unknown function, which is transcriptionally induced in response to KAR treatment in a *KAI2*- and *MAX2*-dependent manner, and currently represents the best-characterized KAR marker gene in Arabidopsis [4, 33]. Despite their similarity, KAI2 and D14 cannot replace each other in Arabidopsis, as shown by promoter swap experiments [34]. This indicates that their expression pattern does not determine their signaling specificity. Instead, this is reached by ligand-receptor preference, and most likely the tissue-specific presence of their ligands, as well as distinctive interaction with other proteins, such as repressors of the SMXL family to trigger downstream signaling [35].

In Arabidopsis and rice, in which KAR/KL signalling has so far been mostly studied, *KAI2* is a single copy gene. However, *KAI2* has multiplied and diversified in other species. For example, the *Physcomitrella patens* genome contains 11 genes encoding KAI2-like proteins [36]. Of these some preferentially bind KAR and others the SL 5-deoxystrigol *in vitro* and this preference is determined by polymorphic amino acids in a loop that determines the rigidity of the ligand-binding pocket [37]. The genomes of parasitic plants of the Orobanchaceae also contain several *KAI2* copies. Some of these have evolved to perceive strigolactones, some can restore KAR-responses in Arabidopsis *kai2* mutants, and others do not mediate responses to any of these molecules in Arabidopsis [12, 38, 39]. Thus, in plant species with an expanded KAI2-family there is scope for a diverse range of ligands and ligand-binding specificities, as well as for diverse protein interaction partners. Apart from discriminating KARs from SLs, it was very recently reported that *KAI2* genes have diversified in the genome of the fire follower *Brassica tournefortii* to encode KAI2 receptors, with different ligand preferences towards KAR_1_ and KAR_2_ [40]. Of these, *Bt*KAI2a mediates stronger responses to KAR_2_, while *Bt*KAI2b mediates stronger responses to KAR_1_, when expressed in the heterologous Arabidopsis background. This binding preference is determined by two valine (*Bt*KAI2a) to leucine (*Bt*KAI2b) substitutions at the ligand binding pocket [40]. Also among plants with a single copy *KAI2* gene, the responsiveness to karrikin molecules can differ significantly: Arabidopsis plants respond more strongly to KAR_2_ than to KAR_1_ [4, 15]. In contrast, rice roots did not display any transcriptional response to KAR_2_, not even for the marker gene *DLK2* [16]. It is yet unclear what determines these differences in KAR_2_ responsiveness among plant species.

Legumes comprise a number of agronomically important crops and they are special among plants as most species in the family can form nitrogen-fixing root nodule symbiosis with rhizobia in addition to arbuscular mycorrhiza. Given the possible diversity in KAI2-ligand specificities among plant species, we characterized the karrikin receptor machinery in a legume, using *L. japonicus* as a model. We found that *KAI2* has duplicated prior to the diversification of legumes and that *L. japonicus* KAI2a and KAI2b differ in their binding preferences to synthetic ligands *in vitro* and in the heterologous Arabidopsis *kai2* and *kai2 d14* mutant backgrounds. We demonstrate that these ligand binding preferences can be explained by substitution of a highly conserved phenylalanine to a tryptophan in the binding pocket of *Lj*KAI2b. This tryptophan occurs rarely also in other unrelated angiosperm species, and seems to have arisen several times independently. Two additional polymorphic residues that are conserved in the KAI2a and KAI2b clades across several legumes may also contribute to ligand binding preference. In addition, we found a surprising organ-specific responsiveness to synthetic KAI2-ligands, with *L. japonicus* hypocotyl development responding to KAR_1_, KAR_2_ and the strigolactone/karrikin analog *rac*-GR24, and root system development responding only to KAR_1_. These responses depend only on *Lj*KAI2a in hypocotyls, while *Lj*KAI2a and *Lj*KAI2b operate redundantly in roots. Together these findings suggest that a diversity of mechanisms may influence KAR/KL responses including receptor-ligand binding specificity or organ-specific interaction of KAI2 with other proteins.

## Results

### *KAI2* underwent duplication prior to diversification of the legumes

To characterize the karrikin and the strigolactone perception machinery in *L. japonicus* we retrieved KAI2, D14 and MAX2 by protein BLAST using Arabidopsis KAI2, D14 and MAX2 as templates. A phylogenetic tree revealed that *LjD14* (*Lj5g3v0310140.4*) is a single copy gene whereas *LjKAI2* has duplicated (Fig 1), resulting in two paralogs *LjKAI2a* (*Lj2g3v1931930.1*) and *LjKAI2b* (*Lj0g3v0117039.1*). The *KAI2* duplication event must have occurred prior to the diversification of the legumes or at least before the separation of the Millettioids and the ‘Hologalegina’ clade [41] because a similar duplication pattern as in *L. japonicus* (Hologalegina) is also detected in pea, *Medicago truncatula* (both Hologalegina) and soybean (Millettioid). The Millettioid soybean genome additionally contains a third, more distantly related *KAI2* copy (*KAI2c*).

**Figure 1.**
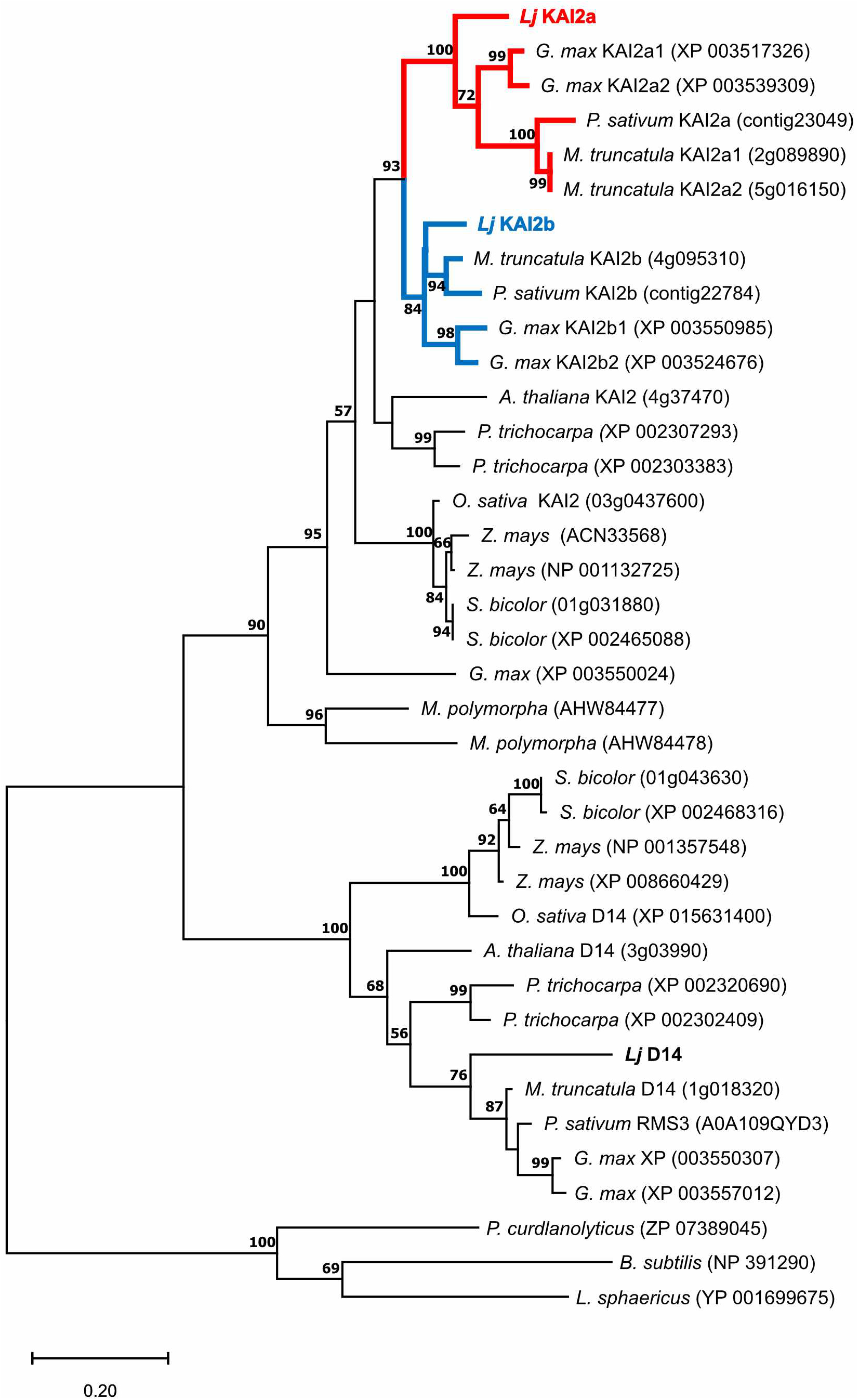
The KAI2 gene underwent duplication prior to diversification of the legumes. Phylogenetic tree of KAI2 and D14 rooted with bacterial RbsQ from indicated species (*Lotus japonicus; Glycine max; Pisum sativum; Medicago truncatula; Arabidopsis thaliana; Populus trichocarpa; Oryza sativa; Zea mays; Sorghum bicolor; Marchantia polymorpha*). MEGAX was used to align the protein sequences with MUSCLE and generate a tree inferred by Maximum Likelihood method [72]. The tree with the highest log likelihood (−7359.19) is shown. The percentage of trees in which the associated taxa clustered together is shown next to the branches. Values below 50 were ignored. *KAI2* duplication in the legumes is highlighted by red and blue branches.

The F-box protein-encoding gene *LjMAX2* also underwent duplication likely as a result of whole genome duplication, because the two *LjMAX2* copies are in two syntenic regions of the genome (S1A Fig). However, only one *LjMAX2* copy (*Lj3g3v2851180.1*) is functional. The other copy *ΨMAX2-like* (*Lj0g3v0059909.1*) appears to be a pseudogene, as it contains an early stop codon, thus encoding a putative truncated protein of 216 instead of 710 amino acids (S1B Fig). It appears that an insertion of one nucleotide into *ΨMAX2-like* created a frameshift, as manual deletion of thymine 453 restores a correct nucleotide and amino acid sequence (S1B Fig).

We hypothesized that *L. japonicus* (and other legumes) retained two intact *KAI2* copies because they may have functionally diverged, perhaps through changes in their expression pattern and/or sequence, possibly resulting in a divergent spatial distribution, ligand affinity and/or ability to interact with other proteins. We examined transcript accumulation of *LjKAI2a* and *LjKAI2b*, as well as *LjD14* and *LjMAX2* in different organs of *L. japonicus* (S2 Fig). Overall, both *LjKAI2a* and *LjKAI2b* transcripts accumulated to higher levels than those of *LjD14* and *LjMAX2. LjKAI2a* transcripts accumulated approximately 100-fold more in aerial organs than *LjKAI2b*, whereas *LjKAI2b* accumulated 10-fold more than *LjKAI2a* in roots of adult plants, which were grown in a sand-vermiculite mix in pots under long day conditions (16h light / 8h dark). However, 1-week-old seedlings grown on water-agar in Petri dishes under short-day conditions (8h light / 16h dark), displayed 10-fold higher transcript levels of *LjKAI2a* than *LjKAI2b* in both roots and hypocotyls (S2B Fig). Thus, *LjKAI2a* and *LjKAI2b* are regulated in an organ-specific, age- and/or environment-dependent manner, suggesting that their individual expression involves at least partially different transcriptional regulators.

Fusions of the four corresponding proteins with T-Sapphire or mOrange in transiently transformed *Nicotiana benthamiana* leaves showed similar subcellular localization as in Arabidopsis and rice [16, 17, 42, 43]. T-Sapphire-MAX2 was detected exclusively in the nucleus, while the α/β-hydrolases (D14, KAI2a and KAI2b) fused to mOrange localized to the nucleus and cytoplasm (S3A Fig). Western blot analysis confirmed that the mOrange signal observed in the cytoplasm resulted from the full-length fusion protein and not from free mOrange (S3B Fig).

### *L. japonicus KAI2a, KAI2b* and *D14* can replace their orthologs in Arabidopsis

To examine whether both *Lj*KAI2a and *Lj*KAI2b function in a canonical manner, we employed a well-established hypocotyl elongation assay in Arabidopsis [12, 34], after transgenically complementing the *Arabidopsis thaliana kai2-2* mutant [4] with *LjKAI2a* and *LjKAI2b* driven by the *AtKAI2* promoter. Both restored inhibition of hypocotyl elongation in the *kai2-2* mutant (Fig 2A). *LjD14* driven by the *AtKAI2* promoter was unable to restore hypocotyl growth inhibition, but it restored repression of shoot branching of the Arabidopsis *d14-1* mutant [4], when driven by the Arabidopsis *D14* promoter. As expected, *LjKAI2a* and *LjKAI2b* could not do the same (Fig 2B and 2C). Together with the phylogenetic analysis (Fig 1), these results demonstrate that *L. japonicus KAI2a* and *KAI2b* are both functional orthologs of the Arabidopsis karrikin/KL receptor gene *KAI2*, whereas *L. japonicus D14* is the functional orthologue of the Arabidopsis strigolactone receptor gene *D14*. Furthermore, the *L. japonicus KAI2* genes are not interchangeable with *D14* [34].

**Figure 2.**
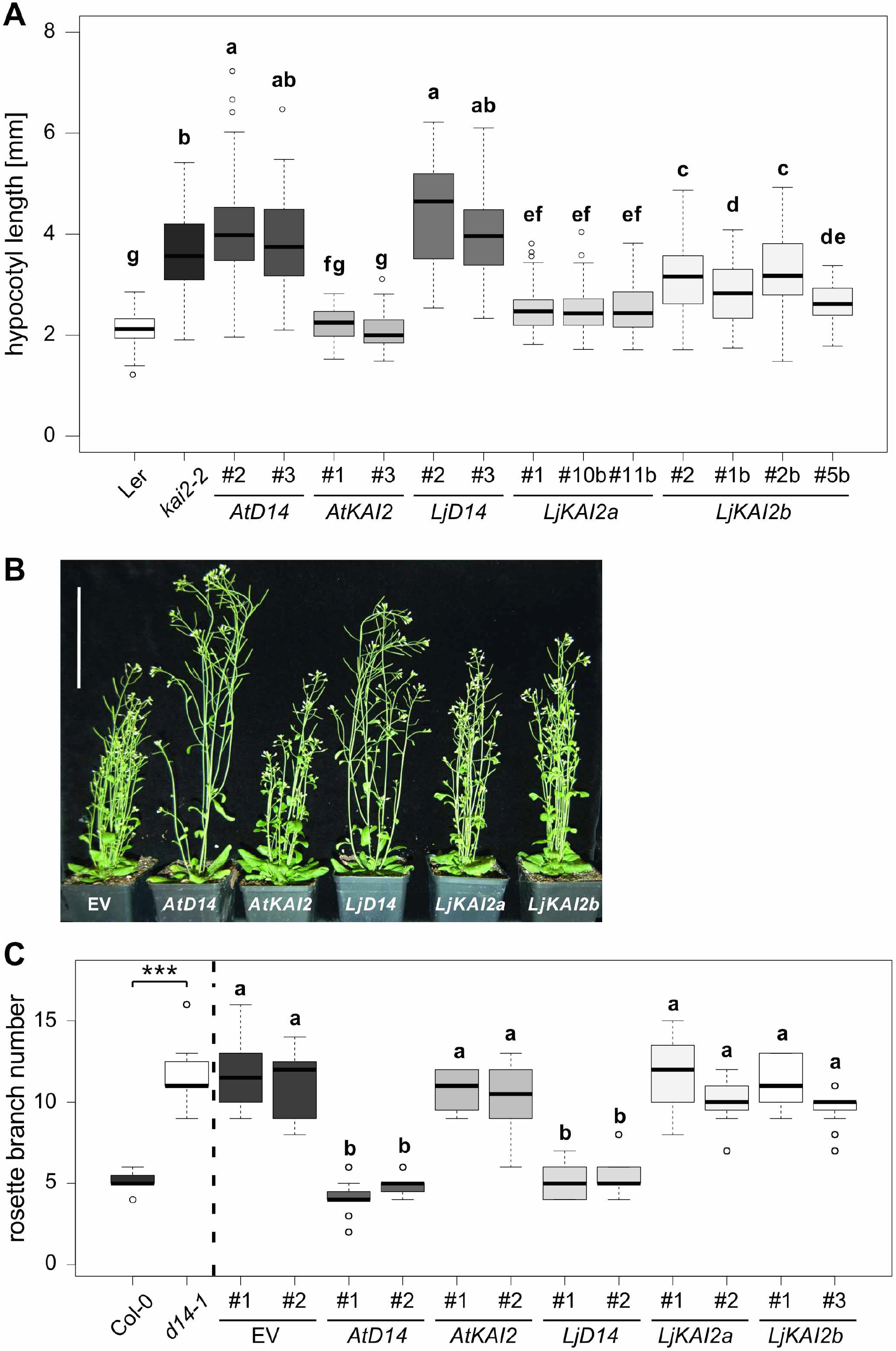
*Lotus japonicus D14, KAI2a* and *KAI2b* can replace *D14* and *KAI2* in Arabidopsis, respectively. (**A**) Hypocotyl length of *A. thaliana* wild-type (L*er*), *kai2-2* and *kai2-2* lines complemented by *AtD14, AtKAI2, LjD14, LjKAI2a* and *LjKAI2b*, driven by the *AtKAI2* promoter at 6 days post germination (dpg). Seedlings were grown in 8h light / 16h dark periods (n=37-122). (**B**) Shoots of *A. thaliana d14-1*, with an empty vector (EV) or complemented with *AtD14, AtKAI2, LjD14, LjKAI2a* and *LjKAI2b*, driven by the *AtD14* promoter at 26 dpg. Scale bar = 10 cm. (**C**) Rosette branch number at 26 dpg of *A. thaliana* wild-type (Col-0), *d14-1* and *d14-1* lines carrying an empty vector (EV) or plasmids containing *AtD14, AtKAI2, LjD14, LjKAI2a* and *LjKAI2b*, driven by the *AtD14* promoter (n=24). Letters indicate different statistical groups (ANOVA, post-hoc Tukey test).

### *Lotus japonicus* KAI2a and KAI2b differ in their ligand binding specificity

To explore whether *L. japonicus* KAI2a and KAI2b can mediate hypocotyl responses to karrikins, we quantified hypocotyl length of the *Atkai2-2* lines transgenically complemented with *LjKAI2a* or *LjKAI2b* after treatment with KAR_1_ and KAR_2_ (Fig 3A and 3B). Two independent lines complemented with *LjKAI2a* displayed a similar reduction in hypocotyl growth in response to both KAR_1_ and KAR_2_. However, the two lines expressing *LjKAI2b* responded more strongly to KAR_1_ than to KAR_2_, contrasting with the common observation, that Arabidopsis hypocotyl growth tends to be more responsive to KAR_2_ [4, 44]. We wondered if the preference towards a specific KAR compound is also observed with KAI2 from other species. To this end, we tested the karrikin response in a line resulting from a cross of the *kai2* mutant *htl-2* with an Arabidopsis line transgenic for the cDNA of the rice *D14L/KAI2* [16]. In contrast to *LjKAI2b, OsD14L/KAI2* mediated a stronger response to KAR_2_ than to KAR_1_ (Fig 3C). Thus, the differential responsiveness of transgenic Arabidopsis lines to KAR_1_ and KAR_2_ does not result from a general incompatibility of a heterologous KAI2 protein with the Arabidopsis background, but suggests different ligand affinities of the transgenic receptors to the karrikins or their possible metabolised products [34, 40].

**Figure 3.**
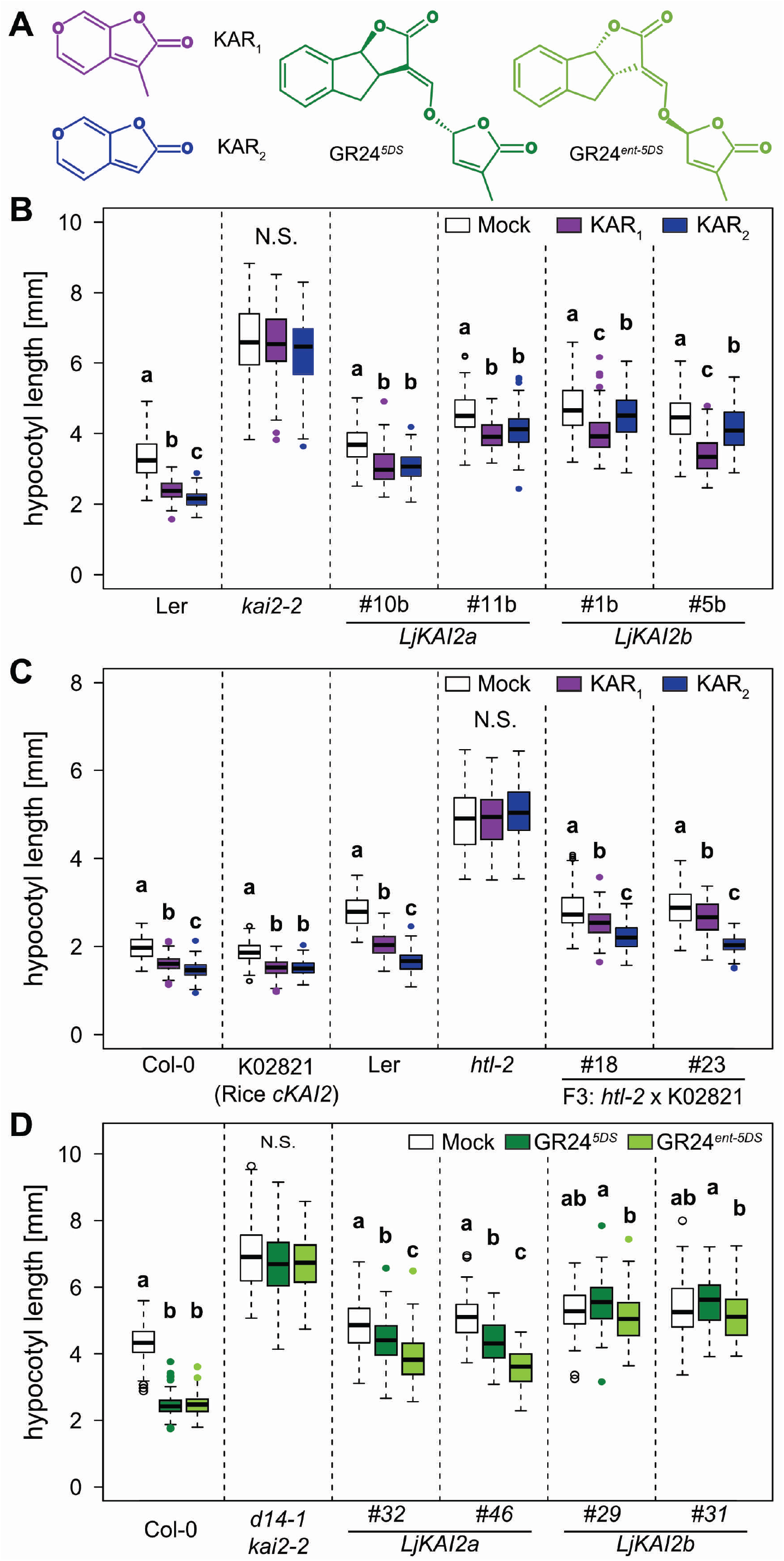
*Lotus japonicus* KAI2a, KAI2b and rice D14L confer divergent hypocotyl growth responses to KAR_1_ and KAR_2_ in Arabidopsis. (**A**) Structures of KAR_1_, KAR_2_, GR24^*5DS*^ and GR24^*ent*-5DS^. (**B-C**) Hypocotyl length of *A. thaliana kai2* mutants complemented with *KAI2* from *A. thaliana*, *L. japonicus* and rice, after treatment with solvent (Mock), 1 μM KAR_1_ or KAR_2_ at 6 dpg. (**B**) L*er* wild-type, *kai2-2* and *kai2-2* lines complemented with *AtKAI2, LjKAI2a* and *LjKAI2b*, driven by the *AtKAI2* promoter (n= 33-128). (**C**) L*er* and Col-0 wild-type, *htl-2* (Ler), K02821-line transgenic for *p35S:OsD14L* (Col-0), and two homozygous F_3_ lines from the *htl-2* x K02821 cross [16] (n= 80-138). (**D**) Hypocotyl length of *A. thaliana* Col-0 wild-type, *d14-1 kai2-2* double mutants, and *d14-1 kai2-2* lines complemented with *LjKAI2a* and *LjKAI2b*, driven by the *AtKAI2* promoter after treatment with solvent (Mock), 1 μM GR24^*5DS*^ or GR24^*ent*-5DS^ (n= 59-134). (**B-D**) Seedlings were grown in 8h light / 16h dark periods. Letters indicate different statistical groups (ANOVA, post-hoc Tukey test).

The two enantiomers of the synthetic strigolactone *rac*-GR24, namely GR24^5DS^ and GR24^*ent*-5DS^, trigger developmental and transcriptional responses via D14 as well as KAI2, respectively, in Arabidopsis [15, 45]. For some KAI2-mediated responses, GR24^*ent*-5DS^ was shown to be more active than karrikin [46] and it has been hypothesized that karrikin may need to be metabolized *in planta*, to yield a high affinity KAI2 ligand, while this may not be nessessary for GR24^*ent*-5DS^ [34, 46, 47]. We examined whether *Lj*KAI2a and *Lj*KAI2b can mediate hypocotyl growth responses to GR24^5DS^ and GR24^*ent*-5DS^ in the *Arabidopsis thaliana d14-1 kai2-2* double mutant background (Fig 3D). Lines expressing *LjKAI2a* responded to both enantiomers with reduced hypocotyl elongation, but displayed a much stronger response to the preferred KAI2 ligand GR24^*ent*-5DS^. Unexpectedly, the lines expressing *LjKAI2b* did not significantly respond to either of the two enantiomers. This contrasting sensitivity to GR24 enantiomers together with the differences in response to KAR_1_ and KAR_2_ suggests that *Lj*KAI2a and *Lj*KAI2b differ in their binding pocket, resulting in divergent affinity to the synthetic ligands.

### Replacement of a conserved phenylalanine by a tryptophan at the binding pocket of KAI2b explains rejection of GR24^*ent*-5DS^

We used differential scanning fluorimetry (DSF) to examine whether purified recombinant *Lj*KAI2a and *Lj*KAI2b (S4A Fig) display a different ligand affinity *in vitro*, employing GR24^5DS^ and GR24^*ent*-5DS^ as model ligands (Fig. 4, S4 Fig). The DSF assay has been widely used for deducing GR24^5DS^ and GR24^*ent*-5DS^ binding to D14 and KAI2 proteins respectively, by means of thermal destabilization [26, 34, 40, 48–50]. However, DSF is unfortunately not suitable for the characterization of karrikin binding [34, 40]. Neither *Lj*KAI2a nor *Lj*KAI2b were destabilized in the presence of GR24^5DS^ (S4B Fig). GR24^*ent*-5DS^ induced a significant thermal destabilization of *Lj*KAI2a at a concentration > 50 μM. In contrast, it did not cause any significant thermal shift of *Lj*KAI2b (Fig 4B), thus recapitulating the difference in hypocotyl growth response between Arabidopsis lines expressing *LjKAI2a* and *LjKAI2b* (Fig 3D).

**Figure 4.**
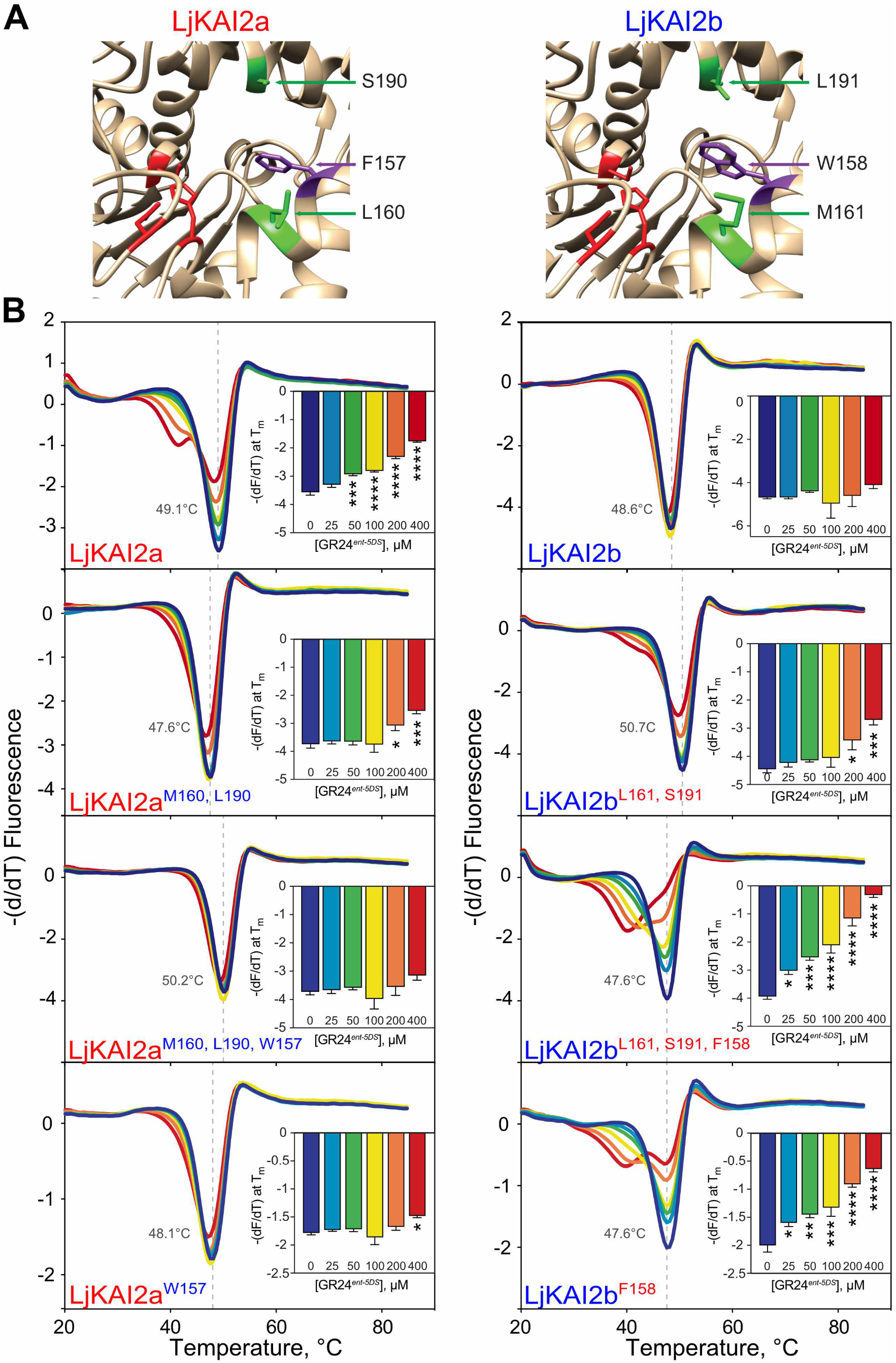
Binding of GR24^*ent*-5DS^ to *Lj*KAI2a is determined by three amino acids. (**A**) The ligand-binding cavity regions of *Lj*KAI2a and *Lj*KAI2b proteins after structural homology modelling on the KAI2 crystal structure of *A. thaliana* [5]. Conserved residues in the cavity that differ between the KAI2a and KAI2b clades, and that are also different between *Lj*KAI2b and *At*KAI2, are shown in green. The phenylalanine residue in *Lj*KAI2a, which is changed to tryptophan in *Lj*KAI2b, is shown in violet. The catalytic triad is coloured in red. (**B**) DSF curves of purified SUMO fusion proteins of wild-type *Lj*KAI2a and *Lj*KAI2b, and versions with swapped amino acids *L/*KAI2a^M160,L190^, *Lj*KAI2b^L161,S191^, *Lj*KAI2a^W157,M160,L190^, *Lj*KAI2b^F158,L161,S191^, *Lj*KAI2a^W157^, *Lj*KAI2b^F158^ at the indicated concentrations of of GR24^*ent*-5DS^. The first derivative of the change of fluorescence was plotted against the temperature. Each curve is the arithmetic mean of three sets of reactions, each comprising four technical replicates. Peaks indicate the protein melting temperature. The shift of the peak in *Lj*KAI2a indicates ligand-induced thermal destabilisation consistent with a protein-ligand interaction. Insets plot the minimum value of (−dF/dT) at the melting point of the protein as determined in the absence of ligand (means ± SE, n = 3). Asterisks indicate significant differences to the solvent control (ANOVA, post-hoc Dunnett test, N.S.>0.05, *≤0.05, **≤0.01, ***≤0.001, ****≤0.0001).

We found 16 conserved amino acid differences between the KAI2a and the KAI2b clade of the investigated legumes (Fig. S5), which may contribute to functional diversification of KAI2a and KAI2b. Modelling of *Lj*KAI2a and *Lj*KAI2b on the KAR_1_-bound *At*KAI2 crystal structure (4JYM) [5] revealed that only three of these were located at the binding pocket, namely L160/M161, S190/L191 and M218/L219 (for the KAI2a/KAI2b comparison, Fig 4A). Out of these three, L219 was conserved between *Lj*KAI2b and the KAI2 proteins from Arabidopsis and rice, which displayed an opposite response pattern to KAR_1_ and KAR_2_ compared to *Lj*KAI2b, when expressed in the Arabidopsis background (Fig. 3). Therefore, we concluded that the M218/L219 polymorphism likely does not play a major role in determining the differential ligand-preference between *Lj*KAI2a and *Lj*KAI2b and discounted it as a candidate. However, we also found that in *Lj*KAI2b exclusively, a highly conserved phenylalanine inside the pocket is replaced by tryptophan at position 158 (S5 Fig). Although this tryptophan is not conserved among other legume KAI2b versions used for the alignment (S5 Fig), we predicted that this bulky residue should have a strong impact on ligand binding.

To understand the impact of the three divergent candidate amino acids on ligand binding, we generated chimeric receptor proteins (Fig 4B). Exchanging only the two amino acids that are conserved across KAI2a and KAI2b clades comprising the investigated legumes (S5 Fig) was sufficient to influence the melting temperature of the two proteins in response to GR24^*ent*-5DS^. *Lj*KAI2a^M160,L190^ became less responsive relative to *Lj*KAI2a and displayed a slight shift in melting temperature only with 200 μM GR24^*ent*-5DS^, whereas *Lj*KAI2b^L161,S191^ gained a weak ability to respond to GR24^*ent*-5DS^ at 200 μM. When all three amino acids were swapped, the melting response to GR24^*ent*-5DS^ was entirely switched between the two receptor proteins: *Lj*KAI2a^M160,L190,W157^ did not display any thermal shift in presence of GR24^*ent*-5DS^, whereas *Lj*KAI2b^L161,S191,F158^ gained a strong response to GR24^*ent*-5DS^ and displayed a thermal shift with ligand concentrations as low as 25 μM. Thus, *Lj*KAI2b^L161,S191,F158^ seemed to be slightly more prone to ligand-induced destabilisation than wild-type *Lj*KAI2a. As W158 appeared to be a critical amino acid for restricting the response to GR24^*ent*-5DS^, we also tested whether swapping F157 with W158 alone would suffice to exchange the ability of the receptors to respond to GR24^*ent*-5DS^. In effect, *Lj*KAI2b^F158^ recapitulated the response of *Lj*KAI2a to GR24^*ent*-5DS^ and and likewise *Lj*KAI2a^W157^ resembled LjKAI2b. Thus, changing this one amino acid in the binding pocket was sufficient to swap ligand specificity. We conclude that the F158/W159 polymorphism predominantly determines the ability of *Lj*KAI2 proteins to bind GR24^*ent*-5DS^, while there is a weaker contribution of L160/M161 and S190/L191.

As an alternative means to probe ligand-receptor interactions, intrinsic tryptophan fluorescence assays confirmed the response of wild-type *Lj*KAI2a and *Lj*KAI2b and all mutant versions to GR24^*ent*-5DS^ (S6 Fig). Unfortunately, because GR24^*ent*-5DS^ precipitated above 500 μM, this assay did not allow us to calculate Kd values because saturation of response could not be achieved. Nevertheless, the qualitative results reiterate the strong impact of F158/W159 on the relative affinities of *Lj*KAI2a and *Lj*KAI2b for GR24^*ent*-5DS^.

To examine whether the three amino acid residues determine ligand discrimination *in planta*, we transformed Arabidopsis *d14 kai2* double mutants with the mutated *LjKAI2a* and *LjKAI2b* genes driven by the Arabidopsis *KAI2* promoter and performed the hypocotyl growth assay in the presence of GR24^*ent*-5DS^. Swapping only the two amino acids conserved in legumes (M160/L161 and S190/L191) was insufficient to exchange the GR24^*ent*-5DS^ response between lines expressing *Lj*KAI2a vs. *Lj*KAI2b. However, swapping all three amino acids, negatively affected the capacity of *Lj*KAI2a^M160,L190,W157^ to mediate a hypocotyl response to GR24^*ent*-5DS^, whereas it reconstituted a response via *Lj*KAI2b^L161,S191,F158^ in three independent transgenic lines (Fig. 5A and 5B). Although these results do not rule out a contribution of M160/L161 and S190/L191 towards ligand preference, they confirm that the phenylalanine to tryptophan substitution at position 157/158 is critical for determining the difference in GR24^*ent*-5DS^ binding preference between the two *L. japonicus* karrikin receptors KAI2a and KAI2b.

**Figure 5.**
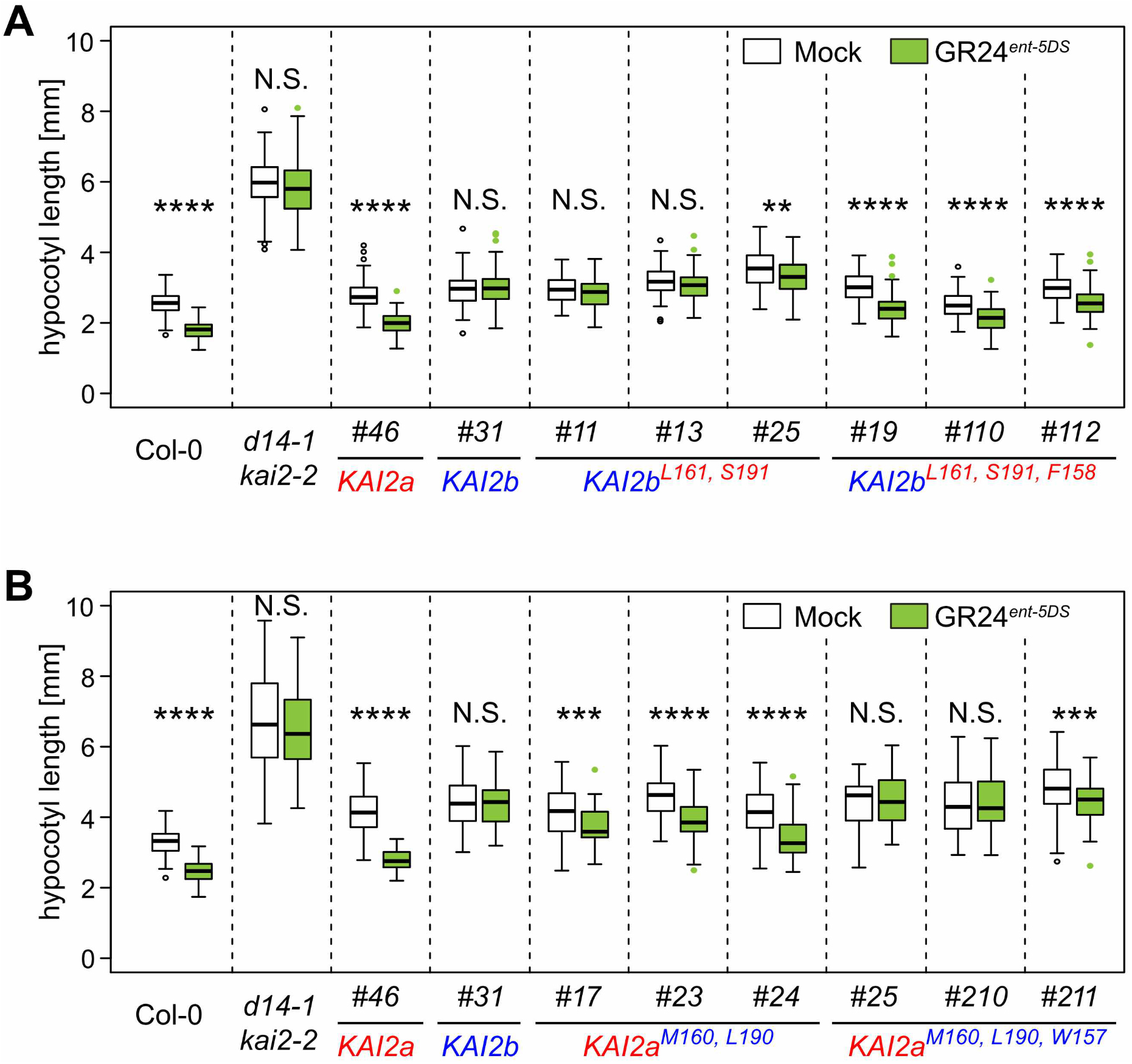
Amino acid swaps reverse sensitivity of *Lj*KAI2a and *Lj*KAI2b to GR24^*ent*-5DS^ in Arabidopsis hypocotyls. Hypocotyl length of *A. thaliana* Col-0 wild-type, *d14-1 kai2-2* double mutants, and *d14-1 kai2-2* lines complemented with *LjKAI2a* and *LjKAI2b* variants driven by the *AtKAI2* promoter and after treatment with solvent (Mock), 1 μM GR24^*5DS*^ or GR24^*ent-5DS*^. (**A**) *LjKAI2a^M160,L190^* and *LjKAI2a^W157,M160,L190^* (n = 46-84). (**B**) *LjKAI2b^L161,S191^* and *LjKAI2b^F158,L161,S191^* (n= 49-102). (**A-B**) Seedlings were grown in 8h light / 16h dark periods Asterisks indicate significant differences versus mock treatment (Welch t.test, *≤0.05, **≤0.01, ***≤0.001, ****≤0.0001).

### The phenylalanine to tryptophan exchange occurred in several unrelated angiosperms independently

The phenylalanine-to-tryptophan transition requires two base changes at position two and three of the codon. We asked whether this change also occurred in other species and searched for KAI2 sequences across the plant phylogeny by BLAST-P against the EnsemblPlants, NCBI and 1KP databases [51], and retrieved KAI2 sequences of the parasitic plants *Striga hermonthica, Orobanche fasciculata* and *Orobanche minor* from Conn et al. 2015 [38]. This analysis showed that the *KAI2c* copy, we detected in the soybean genome (Fig 1) occurred in all analysed genomes of Milletioid legumes, the Genistoid legume *Lupinus albus* and the Mimosoid legume *Prosopsis alba* (S7 Fig). This suggests that the members of the Hologalegina clade, such as *L. japonicus*, have secondarily lost *KAI2c*. Importantly, among the 156 KAI2 sequences we analysed, ten in addition to *Lj*KAI2b contain a tryptophan at the position corresponding to 157 in Arabidopsis KAI2 (S7 Fig). One of them was present in another legume (*Prosopis alba*). Five were present in other eudicots, of which three were in the Lamiales (Paulowniaceae: *Paulownia fargesii;* Phrymaceae: *Erythranthe guttata*, Orobanchaceae: *Orobanche fasciculata*) and two in the Ericales (Primulaceae: *Ardisia evoluta* and *Ardisia humilis*). Furthermore, we found four in monocots of the Bromeliacae (*Ananas comosus*), *Dioscoreaceae* (*Dioscorea rotundata*) and Iridaceae (*Sisyrinchium angustifolium*). In these species, W157 does not co-occur with M160 and L190 as in *Lj*KAI2b, but mostly in combination with the more widely conserved residues L160 and A190 and also with L160 and L190 in *Prosopis alba* and *Ananas comosus* (S7 Fig). The genomes of all dicotyledon species encoding a KAI2 version with W157 contained at least one second copy encoding F157. In the monocot *Dioscorea rotundata* two *KAI2* copies encoded the W157, whereas in *Ananas comosus* and *Sisyrinchium angustifolium*, we detected only one *KAI2* copy. However, we cannot exclude the existence of additional copies as several transcriptomes in the 1KP database are likely incomplete.

In summary, we demonstrate that the F to W transition has occurred several times independently in the angiosperms without co-dependency on M160 and L190 of *Lj*KAI2b, and in most cases it occurred in a duplicate KAI2 version. Thus, the binding pocket of KAI2 proteins appears to be subject to diversification, broadening the range of diverse KAI2-ligand variants that can be recognized, and at the same time extending the opportunities for binding- and signaling-specificity through KAI2 variants with a less (F157) and/or more (W157) restrictive binding-pocket.

### Characterization of *L. japonicus* karrikin and strigolactone receptor mutants

To explore the roles of *LjKAI2a* and *LjKAI2b* in *L. japonicus*, we characterized mutants in these genes as well as in *D14* and *MAX2*. We identified *LORE1* retrotransposon insertions in *L. japonicus KAI2a, KAI2b* and *MAX2* (*kai2a-1, kai2b-3, max2-1, max2-2, max2-3, max2-4*) in available collections [52, 53] and nonsense mutations in *D14* and *KAI2b* (*d14-1, kai2b-1, kai2b-2*) by TILLING [54] (Fig. 6A, S1 Table). Since some of the *max2* and *kai2b* mutants were impaired in seed germination or production (S1 Table) we continued working with *kai2b-1, kai2b-3, max2-3* and *max2-4*. Quantitative RT-PCR analysis revealed that all mutations caused reduced transcript accumulation of the mutated genes in roots of the mutants except for *d14-1* (S8 Fig). Furthermore, the transcript accumulation of *LjKAI2a* and *LjKAI2b* was not affected by mutation of the respective other paralog (S8A Fig).

**Figure 6.**
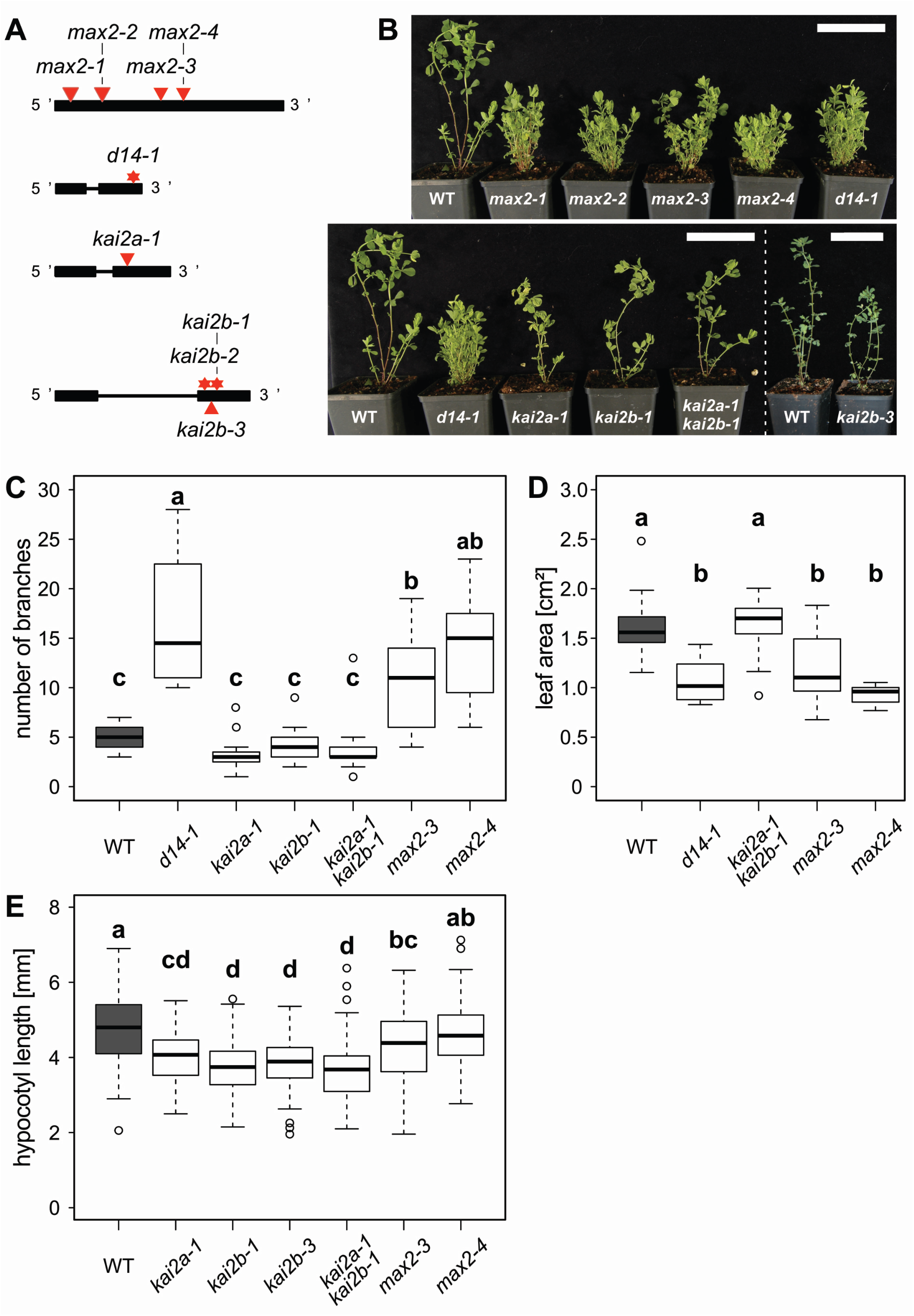
Role of *D14*, *KAI2a, KAI2b* and *MAX2* in shoot and hypocotyl development of *Lotus japonicus*. (**A**) Schematic representation of the *L. japonicus D14, KAI2a, KAI2b* and *MAX2* genes. Black boxes and lines show exons and introns, respectively. *LORE1* insertions are indicated by red triangles and EMS mutations by red stars. (**B**) Shoot phenotype of *L. japonicus* wild-type and karrikin and strigolactone perception mutants at 8 weeks post germination (wpg). Scale bars: 7 cm. (**C**) Number of branches and of *L. japonicus* wild-type, karrikin and strigolactone perception mutants at 7 wpg (n = 12-21). (**D**) Leaf size of the indicated genotypes at 9 wpg (n = 12-15 plants with an average of 3 leaves). (**E**) Hypocotyl length of the indicated genotypes of *L. japonicus* under short day conditions (8h light/ 16h dark) at 1 wpg (n = 79-97). (**C-E**) Letters indicate different statistical groups (ANOVA, post-hoc Tukey test).

The LORE1 insertion in the *kai2a-1* mutant is located close (19 bp) to a splice acceptor site. Since some *LjKAI2a* transcript accumulated in the mutant, we sequenced this residual transcript to examine the possibility that a functional protein could still be made through loss of LORE1 by splicing. We found that indeed a transcript from ATG to stop accumulates in *kai2a-1* but it suffers from mis-splicing leading to a loss of the LORE1 transposon plus 15 bp (from 369 - 383), corresponding to five amino acids (YLNDV) at position 124-128 of the protein (S9A-9B Fig). This amino-acid stretch reaches from a loop at the surface of the protein into the cavity of the binding pocket (S9C Fig). The artificial splice variant did not rescue the Arabidopsis *kai2-2* hypocotyl phenotype, confirming that it is not functional *in planta* and showing that the amino acids 124-YLNDV-128 are essential for *Lj*KAI2a function (S9D Fig).

### Karrikin and *rac*-GR24 cause reduction in hypocotyl growth of *L. japonicus* in an *LjKAI2a*-dependent manner

The *d14-1* and all *max2* mutants displayed increased shoot branching, indicating that the *L. japonicus* strigolactone receptor components D14 and MAX2 are involved in shoot branching inhibition (Fig 6B and 6C), as for Arabidopsis, pea and rice [4, 43, 55, 56]. In addition, *d14* and *max2* mutants had smaller leaves (Fig 6D), a phenotype that has not yet been associated with strigolactone signalling in other dicotyledon species. Surprisingly, *kai2a* and *kai2b* single mutants as well as *kai2a-1 kai2b-1* double mutants or *max2* mutants did not display the canonical elongated hypocotyl phenotype, which is observed in Arabidpsis [4] also in white light conditions (Fig 2, 3 and 5). If anything, the *kai2a-1 kai2b-1* and *max2* mutant hypocotyls were shorter than those of the wild type (Fig. 6E). This indicates that the requirement of KL perception for suppression of hypocotyl elongation under white light is not conserved in *L. japonicus* and/or that KL may not be produced under these growth conditions.

To examine whether *L. japonicus* hypocotyls are responsive to karrikin treatment, we measured the dose-response of hypocotyl growth in wild-type to KAR_1_, KAR_2_ and also to *rac*-GR24. Hypocotyl elongation of wild type plants was progressively inhibited with increasing concentrations of all three compounds (S10A Fig). However, it was not suppressed by KAR_1_ or KAR_2_ treatment in the *kai2a-1 kai2b-1* double mutant and the *max2-4* mutant (S10B-10C Fig). This demonstrates that similar to Arabidopsis, the hypocotyl response to karrikin of *L. japonicus* depends on the KAI2-MAX2 receptor complex. We also examined the KAR_1_ response of *kai2a* and *kai2b* single mutant hypocotyls and found that *kai2a-1* did not significantly respond to KAR_1_ and KAR_2_, while the two allelic *kai2b* mutants showed reduced hypocotyl growth in response to both karrikins (S10B Fig). The transcript accumulation pattern of *DLK2* (*Lj2g3v0765370*) – a classical karrikin marker gene in Arabidopsis [4, 45] – was consistent with this observation and *DLK2* was induced in hypocotyls by KAR_1_ and KAR_2_ in a *LjKAI2a*-dependent but *LjKAI2b*-independent manner (S10D Fig). *rac*-GR24 treatment induced an increase of *DLK2* transcripts in a *LjKAI2b*-independent, partially *LjKAI2a*-dependent, and fully *MAX2*-dependent manner, suggesting that this induction is mediated via *Lj*KAI2a (GR24^*ent*-5DS^) and *Lj*D14 (GR24^5DS^), similar to Arabidopsis [45] (S10C Fig). In summary, *LjKAI2a* appears to be necessary and sufficient to perceive karrikins and GR24^*ent*-5DS^ in the *L. japonicus* hypocotyl, possibly because expression of *LjKAI2b* in hypocotyls is too low under short day conditions (S2B Fig).

### *L. japonicus* root system architecture is modulated by KAR_1_ but not by KAR_2_ treatment

*rac*-GR24 treatment can trigger root system architecture changes in Arabidopsis and *Medicago truncatula* [57–59], and it has recently become clear for Arabidopsis that lateral and adventitious root formation is co-regulated by karrikin and strigolactone signalling [15, 25]. We examined whether *L. japonicus* root systems respond to *rac*-GR24, KAR_1_ and KAR_2_ (Fig. 7A). Surprisingly, in contrast to Arabidopsis and *M. truncatula*, *L. japonicus* root systems responded neither to *rac*-GR24 nor to KAR_2_. Only KAR_1_ treatment led to a dose-dependent decrease in primary root length and an increase of post-embryonic root (PER) number, and thus to a higher PER density (Fig. 7A). PERs include lateral and adventitious roots that can be difficult to distinguish in *L. japonicus* seedlings grown on Petri dishes. The instability of *rac*-GR24 over time in the medium could potentially prevent a developmental response of the root to this compound in our experiments [60]. However, refreshing the medium with new *rac*-GR24 or karrikins at 5 days post-germination, did not alter the outcome (Fig. 7B). Consistently, we observed *DLK2* induction in roots after KAR_1_ but not after KAR_2_ treatment (Fig. 7C).

**Figure 7.**
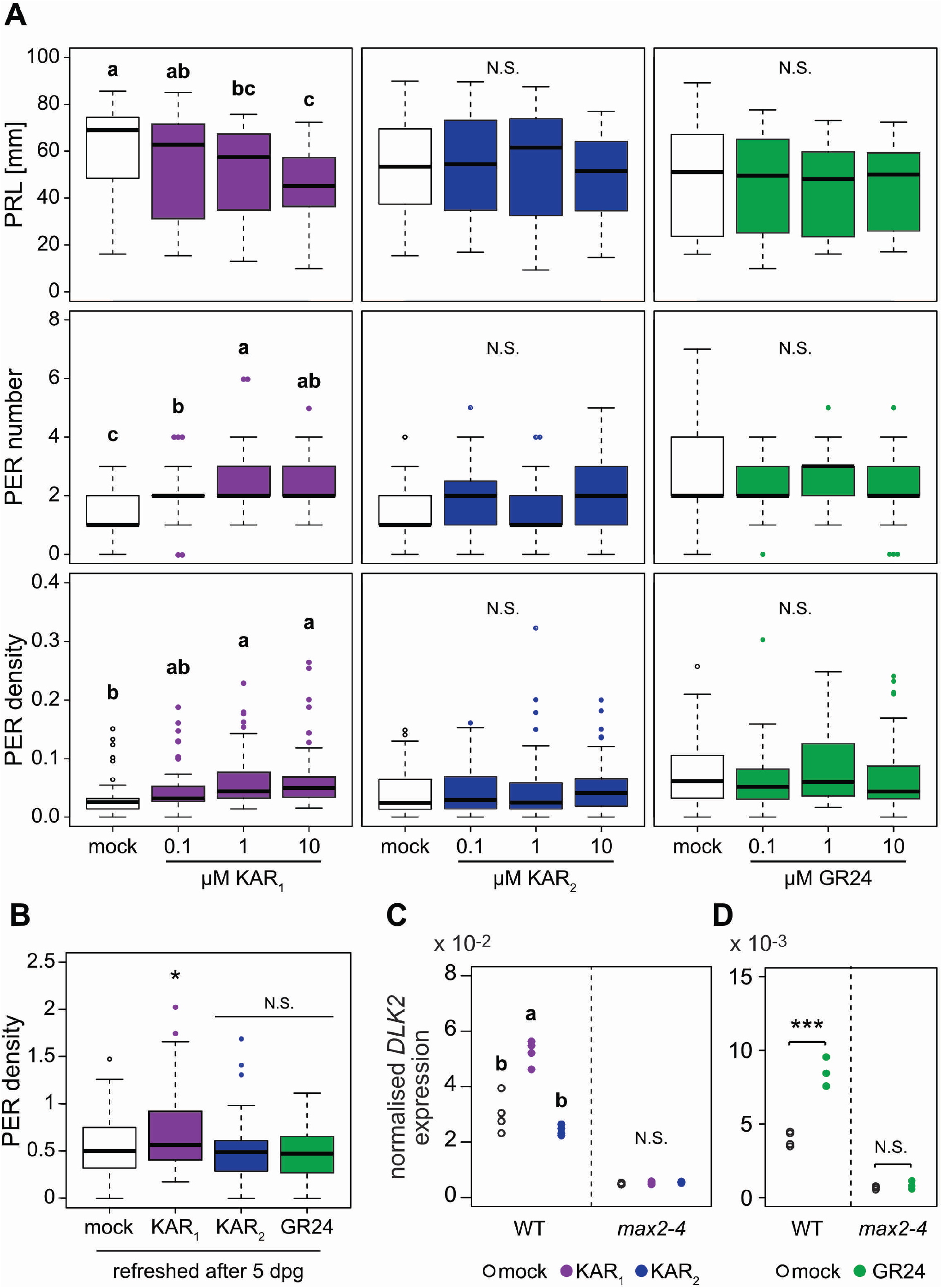
*Lotus japonicus* root system architecture is affected specifically by treatment with KAR_1_ but not KAR_2_. (**A**) Primary root length (PRL), post-embryonic root (PER) number and PER density of wild-type plants 2 wpg after treatment with solvent (M) or three different concentrations of KAR_1_, KAR_2_ or *rac*-GR24 (GR24) (n = 32-57). (**B**) PER density of wild-type plants at 2 wpg and treated with solvent (Mock) 1 μM KAR_1_, 1 μM KAR_2_, or 1 μM *rac*-GR24 (n = 43-51). Plants were transferred onto fresh hormone-containing medium after 5 days. (**C-D**) qRT-PCR-based expression of *DLK2* normalized to *Ubiquitin* expression in roots at 2 wpg after 2 hours treatment with solvent (Mock), (**C**) 1 μM KAR_1_ and 1 μM KAR_2_, (**D**) 1 μM *rac*-GR24 (n = 4). (**A** and **C**) Letters indicate different statistical groups (ANOVA, post-hoc Tukey test). (**B**) Asterisks indicate significant differences (ANOVA, Dunnett test, N.S.>0.05, *≤0.05). (**D**) Asterisk indicate significant differences versus mock treatment (Welch t.test, *≤0.05, **≤0.01, ***≤0.001).

Together with the hypocotyl responses to KAR_1_, KAR_2_ and *rac*-GR24 this indicates organ-specific sensitivity or responsiveness to these three compounds in *L. japonicus* with a more stringent uptake, perception and/or response system in the root.

Surprisingly, we found that the roots responded to *rac*-GR24 treatment with increased *DLK2* transcript accumulation (Fig. 7D) although no change in root architecture was observed in response to this treatment (Fig. 7A). This suggests that different ligands may be transported to different tissues or may have a divergent impact on receptor conformation, thereby mediating different downstream responses. To confirm the contrasting responses of *L. japonicus* root systems to KAR_1_ and *rac*-GR24, and to test whether they result from divergent molecular outputs, we examined transcriptional changes after one, two and six hours treatment of *L. japonicus* wild-type roots with KAR_1_ and *rac*-GR24 using microarrays. Statistical analysis revealed a total number of 629 differentially expressed (DE) genes for KAR_1_-treated and 232 genes for *rac*-GR24-treated roots (Table S2). In agreement with previous reports from Arabidopsis and tomato [44, 61, 62] the magnitude of differential expression was low. Most of the DE genes upon KAR_1_ and *rac*-GR24 treatment responded solely after 2h (S11 Fig). Interestingly, only a minority of 48 genes responded in the same direction in response to both KAR_1_ and *rac*-GR24, while the majority of genes responded specifically to KAR_1_ (580 DEGs) or *rac*-GR24 (169 DEGs). If *rac*-GR24 were to simply mimic the effect of KAR (GR24^*ent*-5DS^) and SL (GR24^5DS^) on roots, one would have expected a large overlap with KAR_1_ responses, and in addition a number of non-overlaping DEGs, regulated through D14. In summary, the microarray experiment confirmed largely non-overlapping responses of *L. japonicus* root response to KAR_1_ and *rac*-GR24.

### Both *Lj*KAI2a and *Lj*KAI2b mediate root architecture-responses to KAR_1_ but only *Lj*KAI2a mediates *DLK2* expression in response to GR24^*ent*-5DS^

To inspect which α/β-hydrolase receptor mediates the changes in *L. japonicus* root system architecture in response to KAR_1_ treatment, we examined PER density in the karrikin receptor mutants. The *Ljkai2a-1 kai2b-1* double mutant and the *max2-4* mutant did not respond to KAR_1_ treatment with changes in root system architecture (Fig. 8A, S12 Fig). With 1 μM KAR_1_, we obtained contradictory results for the single *kai2a* and *kai2b* mutants in independent experiments (S12A and S12C Fig). However, *kai2a* and *kai2b* single mutants but not the *kai2a kai2b* double mutant responded to a slightly higher concentration of 3 μM KAR_1_ (Fig. 8A), indicating that *Lj*KAI2a and *Lj*KAI2b redundantly perceive KAR_1_ (or a metabolite thereof) in *L. japonicus* roots. This pattern was mirrored by *DLK2* expression in roots: both *kai2a* and *kai2b* single mutants responded to KAR_1_ with increased *DLK2* expression, while the *kai2a-1 ka2b-1* double mutant and the *max2-4* mutant did not respond (Fig. 8B). Since *Lj*KAI2b did not respond to GR24^*ent*-5DS^ *in vitro* as well as in the heterologous Arabidopsis background (Fig 4 and 5, S6 Fig), we examined its ability to mediate *DLK2* induction by GR24^*ent*-5DS^ in *L. japonicus* roots. Wild-type and *kai2b* mutant roots responded to GR24^*ent*-5DS^ with increased *DLK2* expression, but this was not the case for *kai2a* roots confirming that *Lj*KAI2b cannot bind and mediate responses to GR24^*ent*-5DS^ (Fig 8C). In summary, *Lj*KAI2a and *Lj*KAI2b act redundantly in roots in mediating responses to KAR_1_ but only KAI2a can perceive GR24^*ent*-5DS^.

**Figure 8.**
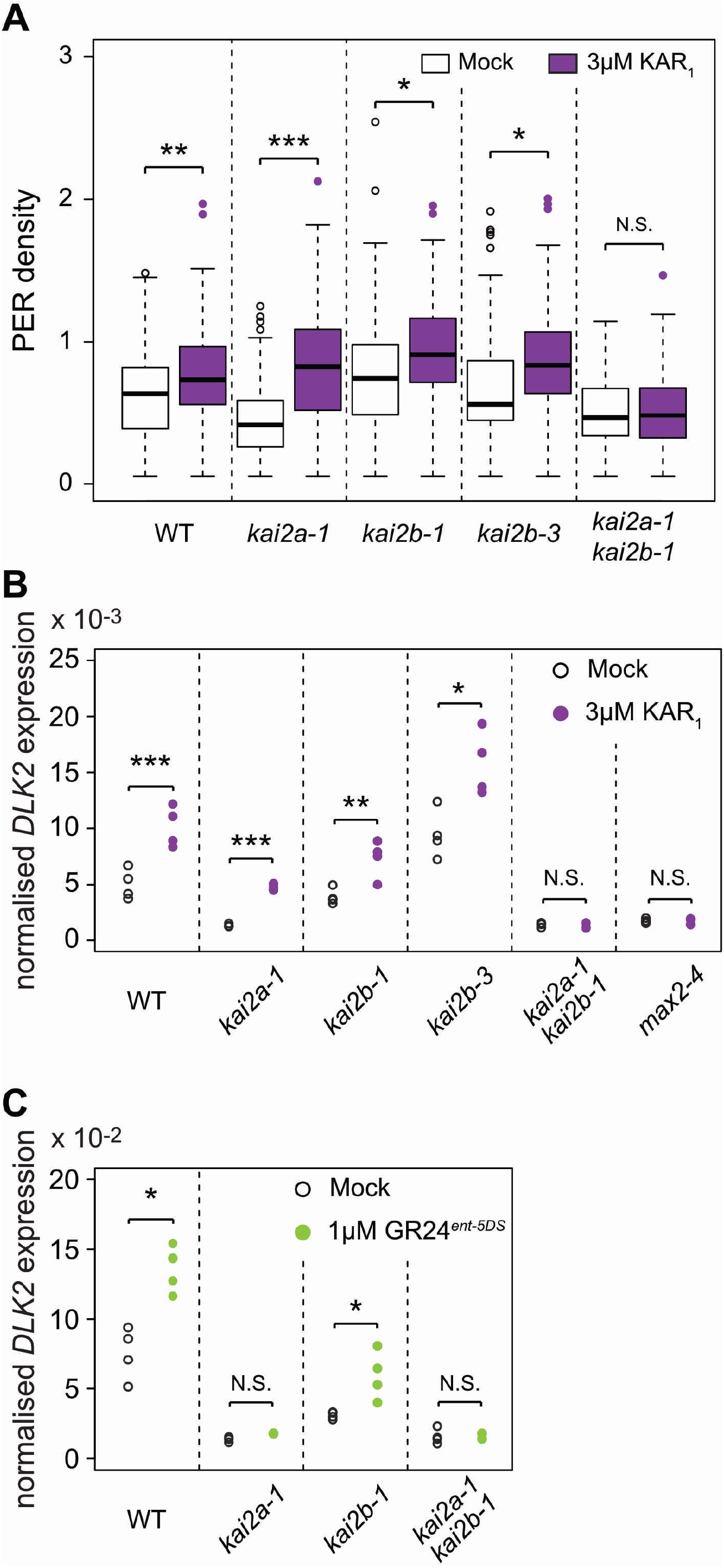
*Lj*KAI2a and *Lj*KAI2b operate redundantly in the response of roots to KAR_1_. (**A**) Post-embryonic-root (PER) density of *L. japonicus* plants, 2 wpg after treatment with solvent (M) or 3 μM KAR_1_ (n=34-72). (**B-C**) qRT-PCR-based expression of *DLK2* in roots of *L. japonicus* plants at 2 wpg after 2 hours treatment with solvent (Mock) or **(B)** 3 μM KAR_1_ or **(C)** 1μM GR24^*ent-5DS*^. Expression values were normalized to those of the housekeeping gene *Ubiquitin* (n= 3-4). (**A-C**) Asterisks indicate significant differences versus mock treatment (Welch t.test, *≤0.05, **≤0.01, ***≤0.001).

## Discussion

Gene duplication followed by sub- or neofunctionalization is an important driver in the evolution of complex signalling networks and signalling specificities during the adaptation to new or diverse environments. In the legumes, the karrikin receptor gene *KAI2* multiplied possibly during the whole genome duplication that occurred in the Papilionoidaea before the diversification of legumes 59 million years ago [63]. While the Mimosoids, Genistoids and Milletioids, contain three different KAI2 versions, the Hologalegina clade appears to have lost one of them, retaining the more closely related *KAI2a* and *KAI2b* versions. Here, we provide evidence that *L. japonicus* KAI2a and KAI2b diversified in their ligand-binding specificity as well as organ-specific requirement (Fig 9).

**Figure 9.**
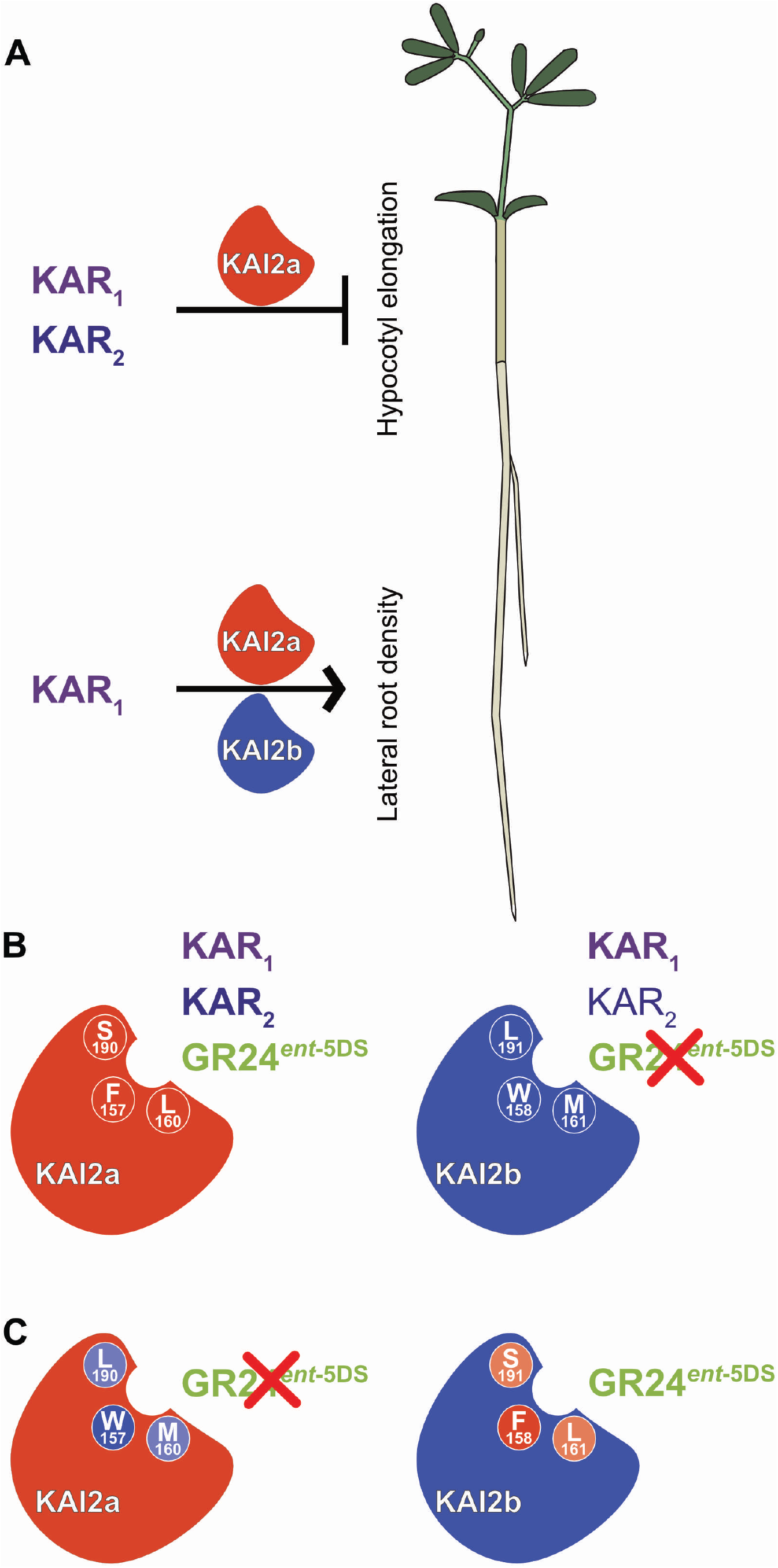
*L. japonicus* KAI2a and KAI2b display organ-specific redundancy and differ in their ligand-binding specificity. (**A**) *Lj*KAI2a is required to mediate inhibition of hypocotyl growth in response to KAR_1_ and KAR_2_. In roots *Lj*KAI2a and *Lj*KAI2b redundantly promote lateral root density, but only in response to KAR_1_ treatment. (**B**) Upper panel: In the Arabidopsis *kai2-2* background *Lj*KAI2a mediates hypocotyl growth inhibition in response to KAR_1_, KAR_2_ and GR24^*ent*-5DS^. In the same background, *Lj*KAI2b mediates a stronger response to KAR_1_ than to KAR_2_ and no response to GR24^*ent*-5DS^ (indicated by a red cross). Three divergent amino acids at the binding pocket are indicated in white. Lower panel: Swapping the three divergent amino acids in the binding pocket reconstitutes GR24^*ent*-5DS^ activity through *Lj*KAI2b and abolishes GR24^*ent*-5DS^ activity through *Lj*KAI2a. Among the three amino acids F157/W158 are decisive for GR24^*ent*-5DS^ binding (strong colors), while L160/M161 and S190/L191 play a weaker role (pale colors). Amino acids from *Lj*KAI2a have a red/pale red and amino acids from *Lj*KAI2b a violet/pale background.

*Lj*KAI2a and *Lj*KAI2b differ in their quantitative sensitivity to KAR_1_ and KAR_2_, which vary only by the presence of one methyl group in KAR_1_ (Fig. 3B). An increased hydrophobicity of the *Lj*KAI2b binding pocket as compared to *Lj*KAI2a may mediate the preference towards the more hydrophobic KAR_1_, similar to the fire-following plant *Brassica tournefortii* [40]. The difference in ligand preference of *Lj*KAI2a vs. *Lj*KAI2b is more dramatic for GR24^*ent*-5DS^, an enantiomer of the synthetic strigolactone analogue *rac*-GR24, which acts through Arabidopsis KAI2 when applied to plants, promotes interaction of KAI2 with SMAX1 in yeast and binds to *At*KAI2 *in vitro* [8, 15, 34, 45, 46]. We show that *Lj*KAI2a mediates strong hypocotyl growth responses to GR24^*ent*-5DS^ in Arabidopsis as well as transcriptional activation of *DLK2* in *Lotus japonicus* roots. *Lj*KAI2b is incapable of triggering these responses to the compound, while being able to induce the same responses upon KAR_1_ treatment. The dramatic difference in the ability of *Lj*KAI2a and *Lj*KAI2b to bind GR24^*ent*-5DS^ is confirmed *in vitro* by DSF and intrinsic tryptophan fluorescence assays. Together, these results demonstrate that the individual α/β-fold hydrolase receptor is sufficient to explain ligand sensitivity in planta.

Identifying the determinants of ligand-binding specificity of D14 and different KAI2 and KAI2-like proteins is an area of active research. Binding specificity of D14 and KAI2 to SLs and KARs respectively has been associated with the geometry and size of the binding pocket [6, 64]. Changes in amino acid residues located in the pocket of divergent KAI2 versions in parasitic weeds have enabled alterations in pocket architecture and evolution of a chimeric receptor that perceives strigolactones like D14 but mediates germination like KAI2 [38, 39]. The rigidity of lid helices forming the tunnel of the binding pocket have been proposed to determine specificity of KAI2-like proteins for strigolactone-like molecules vs. KAR_1_ in *Physcomitrella patens* [37].

We identified three amino acids at the ligand-binding pocket that differ between *Lj*KAI2a and *Lj*KAI2b. Two of these are conserved across the legume KAI2a and KAI2b clades, namely L160 and S190 in KAI2a and M161 and L191 in KAI2b. This pattern of conservation suggests functional relevance in maintaining flexibility for different KAI2 ligands in legumes. Indeed, exchanging these two amino acids slightly changes the thermal instability of the two KAI2 versions in the DSF assay. Neither amino acid change is predicted to substantially impact the pocket volume or geometry but the amino acids of *Lj*KAI2b are more hydrophobic, which may explain the preference for the more hydrophobic KAR_1_ over KAR_2_. A similar phenomenon was observed in *Brassica tournefortii*, a fire-following weed that has two functional *KAI2* genes [40]. Similar to the situation in *L. japonicus, Bt*KAI2b mediated a greater sensitivity to KAR_1_ over KAR_2_ in the Arabidopsis background, while it was the reverse for *Bt*KAI2a. This was explained by one amino acid polymorphism in the binding pocket towards a more hydrophobic amino acid (V98L) in *Bt*KAI2b. Notably, this residue (V98L) is in a very different position than the polymorphics residues in *L. japonicus* KAI2a/KAI2b, suggesting that the receptors are highly plastic and that similar binding-specificities may be achieved by changing hydrophobicity in different positions of the pocket.

Exchanging L160/M161 and S190/L191 between *L. japonicus* KAI2a and KAI2b was sufficient to change their sensitivity to GR24^*ent*-5DS^ in the DSF *in vitro* assay. However, the developmental response of Arabidopsis hypocotyls was hardly changed, possibly because *in vivo*, suboptimal ligand binding to the receptor can be stabilized by interacting proteins. A third amino acid difference (F157/W158) between the two KAI2 proteins occurs in *L. japonicus*. This residue critically determines sensitivity to GR24^*ent*-5DS^ *in vitro* as well as in Arabidopsis likely because the bulky tryptophan may sterically hinder GR24^*ent*-5DS^ binding, while still allowing binding of the smaller karrikins. In fact, swapping F157 with W158 alone was sufficient to swap the ability to respond to GR24^*ent*-5DS^ in DSF as well as intrinsic fluorescence assays.

*B. tournefortii* is a fire-following plant, the seeds of which respond to karrikins with dormancy breaking and germination [40]. Therefore, to maintain two copies of KAI2, one of which is specialized for KAR_1_, the most abundant KAR in smoke, and the other of which may be specialized for the endogenously produced ligand makes adaptive sense for *B. tournefortii*. For *L. japonicus*, which is not a fire-follower, KAR_1_, KAR_2_ and GR24^*ent*-5DS^ are likely not natural KAI2 ligands. Nevertheless, the maintenance of two *KAI2* genes in the Hologalegina legumes, each with amino acid polymorphisms confering differences in binding preferences to artificial ligands, requires an adaptive basis. One possibility is that *L. japonicus* KAI2a and KAI2b have specialized to bind different ligands *in planta*, indicative of legumes producing at least two different versions of the as-yet-unknown KAI2 ligand. The distinct expression patterns and developmental roles of *Lj*KAI2a and *Lj*KAI2b might also be consistent with a tissue-specific diversity of ligands, or even an endogenous ligand versus an exogenous ligand derived from the rhizosphere. From our assays with artificial ligands we extrapolate that *Lj*KAI2b has a higher ligand selectivity than *Lj*KAI2a. The additional amino acid change that occurred in *L. japonicus* but not in the other examined legumes may indicate that the KL bouquet of *L. japonicus* has further diversified. Alternatively, the F to W substitution in *Lj*KAI2b may confer resistance to (a) toxic allelochemical(s) that may be released into the rhizosphere by competing neighbouring plants or by microorganisms. This speculative hypothesis is consistent with the role of *Lj*KAI2b in roots (but not in hypocotyls) and with the observation that the F157W exchange occurred in several unrelated plant species independently that may all encounter compounds capable of blocking KAI2a in their natural habitat. It will be exciting to investigate the biological significance of this receptor sub-functionalization and the putative diversity of their ligands, once the molecule class of KL and its variants have been identified.

In addition to ligand-binding specificity at the level of the receptor, we identified a surprising organ-specific responsiveness to synthetic KAI2 ligands in *L. japonicus*. While hypocotyl growth is inhibited in response to KAR_1_, KAR_2_ and *rac*-GR24, root systems only respond to KAR_1_ with architectural changes (Fig. 9A). To our knowledge such an organ-specific discrimination of different but very similar KAR molecules has not previously so clearly been observed. However, a similar scenario could be at play in rice, in which transcriptome analysis of KAR_2_-treated rice roots identified no differentially expressed gene [16], whereas rice mesocotyls respond with growth inhibition to the same treatment [7, 16]. Although KAI2 can be shown to bind KAR_1_ *in vitro* by isothermal tritration calorimetry or fluorescent microdialysis [5, 6, 65], there is evidence suggesting that KARs are not directly bound by KAI2 *in vivo*, but may be metabolized first to yield the correct KAI2-ligand, which may bind with higher affinity [34, 46]. It is possible that substrate specificities differ among enzymes involved in KAR metabolism in hypocotyls vs. roots. This would imply that the single methyl group, which distinguishes KAR_1_ from KAR_2_, is sufficient to impact specialized metabolism of karrikins. Alternatively, the transport of KAR_2_ or the KAR_2_-derived metabolic product could be limited in the root system, or KAR_2_-derivatives may be rapidly catabolised in roots, thus limiting their effect. While KAR_2_ fails to induce increased PER density as well as *DLK2* expression in *L. japonicus* roots, GR24^*ent*-5DS^ triggers *DLK2* transcript accumulation albeit being unable to increase PER density. *DLK2* induction by GR24^*ent*-5DS^ requires KAI2a, thus involvement of D14 can be excluded. Furthermore, *Lj*KAI2a and *Lj*KAI2b act redundantly in mediating KAR_1_-induced root system changes, excluding the possibility that they are regulated exclusively by *Lj*KAI2b, which cannot bind GR24^*ent*-5DS^. It is tempting to speculate that conformational changes of KAI2 proteins may differ depending on the ligand and that the extend of the change may influence the interaction strength with the karrikin signaling repressor SMAX1, MAX2 and/or additional proteins [8, 66]. Perhaps *DLK2* expression is more sensitive to quantitative SMAX1 removal than genes required to be induced for root system changes, such as the ethylene biosynthesis gene *ACS7* [66]. Alternatively, SMAX1 proteins inhibiting *DLK2* expression are more accessible to the receptor complex, thereby allowing interaction even when the receptor binds a suboptimal ligand, as compared to SMAX1 individuals supressing transcriptional acitivity of genes involved in root system changes; or GR24^*ent*-5DS^ is only taken up into a subset cells, in which SMAX1 removal does not mediate root system changes.

We observed that KAR_1_ treatment triggers increased PER density in *L. japonicus*. This is somewhat contradictory to *kai2* and *max2* mutants in Arabidopsis, which display an increased lateral root density [15]. The discrepancy may result from different physiological optima between the two species or from nutrient conditions in the two experimental systems. We observed the KAR_1_ response of *L. japonicus* root systems in half-Hoagland solution with low phosphate levels (2.5μM PO_4_^3-^) and without sucrose, whereas the root assay in Arabidopsis was conducted in ATS medium (*Arabidopsis thaliana* salts) with 1% sucrose [15]. Phosphate and sucrose levels have previously been described to influence the effect of strigolactone and *rac*-GR24 on Arabidopsis root architecture [57, 67, 68].

In Arabidopsis and rice, KAI2/D14L is required to inhibit hypocotyl and mesocotyl elongation, respectively [3, 4, 16]. Since these two species are evolutionarily distant from each other, but have both retained a function of KL signalling in inhibiting the growth of similar organs, it seemed likely that this function would be conserved among a large number of plant species. Surprisingly, in *L. japonicus*, we observed no elongated hypocotyl phenotype for the *kai2a-1 kai2b-1* double and two allelic *max2* mutants (Fig. 5). However, we could trigger a reduction of hypocotyl elongation by treatment with KAR_1_, KAR_2_ and *rac*-GR24 in the wild type and in a *LjKAI2a* and *LjMAX2*-dependent manner. Perhaps the endogenous KL levels in *L. japonicus* hypocotyls are insufficient to cause inhibition of hypocotyl elongation, at least under our growth conditions.

In summary, we have demonstrated sub-functionalization of two KAI2 copies in *L. japonicus* with regard to their ligand-binding specificity and organ-specific relevance. Furthermore, we find organ-specific responsiveness of *L. japonicus* to two artificial KAI2 ligands. A phenylalanine to tryptophan transition independently occurred in the KAI2-binding pocket in several angiosperms, while a leucine-to-methionine and a serine-to-leucine exchange are conserved in KAI2a and KAI2b across legumes. This conservation and independent multiple occurence of specific amino acid polymorphisms suggests that they bear functional relevance for discriminating diverse KAI2 ligands. Our findings open novel research avenues towards understanding the diversity in KL ligand-receptor relationships and in developmental responses to, as yet, unknown natural as well as synthetic butenolides that influence diverse aspects of plant development.

## Materials and methods

### Plant material and seed germination

The *A. thaliana kai2-2* (Ler background) and *d14-1* (Col-0 background) mutants are from [4], the *d14-1 kai2-2* double mutant from [45], the *htl-2* mutant was provided by Min Ni [69] and the cross with K02821 is from [16]. Seeds were surface sterilized with 70% EtOH. For synchronizing the germination, seeds were placed on ½ MS 1 % agar medium and maintained at 4°C in the dark for 72 hours.

The *L. japonicus* Gifu *max2-1, max2-2, max2-3, max2-4, kai2a-1* and *kai2b-3* mutations are caused by a LORE1 insertion. Segregating seed stocks for each insertion were obtained from the Lotus Base (https://lotus.au.dk, [70]) or Makoto Hayashi (NIAS, Tsukuba, Japan, [53] for *max2-2)*. The *d14-1, kai2b-1* and *kai2b-2* mutants were obtained by TILLING [54] at RevGenUK (https://www.jic.ac.uk/technologies/genomic-services/revgenuk-tilling-reverse-genetics/). Homozygous mutants were identified by PCR using primers indicated in S3 Table. For germination, *L. japonicus* seeds were manually scarified with sand-paper and surface sterilized with 1% NaClO. Imbibed seeds were germinated on 1/2 Hoagland medium containing 2.5μM PO_4_^3-^ and 0.4% Gelrite (www.duchefa-biochemie.com), at 24°C for 3 days in the dark, or on ½ MS 0.8% agar at 4°C for 3 days in dark (only for the experiment in Fig. 6E).

### Phylogenetic, synteny and protein sequence analysis

*Lotus japonicus KAI2, D14* and *MAX2* sequences were retrieved using tBLASTn with *AtKAI2, AtD14* and *AtMAX2*, against the NCBI database, the plantGDB database and the *L. japonicus* genome V2.5 (http://www.kazusa.or.ip/lotus). The presence of MAX2-like was identified by tBLASTn in an in-house genome generated by next generation sequencing using CLC Main Workbench [71]. Pea sequences were found by BLASTn on “pisum sativum v2” database with AtKAI2 as query (https://www.coolseasonfoodlegume.org). For Fig 1, the MUSCLE alignment of the protein sequences was used to generate Maximum-likelihood tree with 1000 bootstrap replicates in MEGAX [72]. For the synteny analysis of *MAX2* and *MAX2-like*, flanking sequences were retrieved from the same in-house genome [71]. For S7 Fig, KAI2 sequences across the plant phylogeny were retrieved by BLAST-P search against the EnsemblPlants, NCBI and 1KP databases [51], in addition, KAI2 sequences of the parasitic plants *Striga hermonthica* and *Orobanche cumana* were retrieved from Conn et al. 2015 [38]. The MUSCLE alignment, generated in MEGAX [72], was used to produce a tree with 1000 bootstrap replicates with IQTREE [73].

### Structural homology modelling of proteins

Proteins were modelled using SWISS-MODEL tool (https://swissmodel.expasy.org) with the *A. thaliana* KAI2 (4JYM) templates [5].

### Bacterial protein expression and purification

Full-length *L. japonicus* coding sequences were cloned into pE-SUMO Amp using primers in S3 Table. Clones were sequence-verified and transformed into Rosetta DE3 pLysS cells (Novagen). Subsequent protein expression and purification were performed as described previously [34], with the following modifications: the lysis and column wash buffers contained 10 mM imidazole, and a cobalt-charged affinity resin was used (TALON, Takara Bio).

### Differential scanning fluorimetry

DSF assays were performed as described previously [34]. Assays were performed in 384-well format on a Roche LightCycler 480 II with excitation 498 nm and emission 640 nm (SYPRO Tangerine dye peak excitation at 490 nm). Raw fluorescence values were transformed by calculating the first derivative of fluorescence over temperature. These data were then imported into GraphPad Prism 8.0 software for plotting. Data presented are the mean of three super-replicates from the same protein batch; each super-replicate comprised four technical replicates at each ligand concentration. Experiments were performed at least twice.

### Intrinsic tryptophan fluorescence (ITF) assay

The ITF assay was performed in 384-well format on a BMG Labtech CLARIOstar multimode plate reader, using black FLUOtrac microplates (Greiner 781076). Reactions (20 μL) were set up in quadruplicate and contained 10 μM protein, 20 mM HEPES pH 7.5, 150 mM NaCl, 1.25% (v/v) glycerol, and 0-500 μM ligand. Ligands were initially prepared in DMSO at 20x concentration, and therefore reactions also contained 5% (v/v) DMSO. Ligands were dissolved in buffer (20 mM HEPES pH 7.5, 150 mM NaCl, 1.25% (v/v) glycerol) at 2× concentration immediately before use, of which 10 μL per well was dispensed with a multichannel pipette. An equivalent volume of 2x solution of protein in buffer was prepared and then dispensed onto the plate using an Eppendorf Multipette with a 0.1 mL tip. The plate was mixed at 120 rpm for 2 min, centrifuged at 500x *g* for 2 min, and then incubated in the dark for 20 min at room temperature. Fluorescence measurements were taken first with fixed wavelength filters (excitation 295/10 nm; longpass dichroic 325 nm; emission 360/20 nm), followed by the linear variable filter monochromator for emission wavelength scans (excitation 295/10 nm, emission 334-400 nm, step width 2 nm, emission bandwidth 8 nm). Measurements were performed at 25 °C using 17 flashes per well for fixed filters or 20 flashes per well for wavelength scans. Gain and focus settings were set empirically for each experimental run. Data were blank-corrected by subtraction of fluorescence values from an identical set of wells containing ligand and buffer but no protein. Data analysis was performed in Graphpad Prism v8.4. Best fit curves were generated from untransformed fluorescence readings using nonlinear regression and the in-built “One site - Total” model, with least squares regression as a fitting method and an asymmetrical (profile-likelihood) 95% confidence interval. As saturation was not reached, only ambiguous values for Kd were returned. For emission wavelength scans, fluorescence values at each wavelength were normalised by expressing as a percentage of the corresponding value from samples lacking ligand.

### Plasmid generation

Genes and promoter regions were amplified using Phusion PCR according to standard protocols and using primers indicated in S3 Table. Plasmids were constructed by Golden Gate cloning [74] as indicated in S4 Table.

### Plant transformation

*kai2-2* and *d14-1* mutants were transformed by floral dip in *Agrobacterium tumefaciens* AGL1 suspension. Transgenic seedlings were selected by mCherry fluorescence and resistance to 20 μg/mL hygromycin B in growth medium. Experiments were performed using T2 or T3 generations, with transformed plants validated by mCherry fluorescence.

### Shoot branching assay

*A. thaliana* and *L. japonicus* were grown for 4 and 7 weeks, respectively in soil in the greenhouse at 16h/8h light/dark cycles. Branches with length >1cm were counted, and the height of each plant was measured.

### Hypocotyl elongation assay

*A. thaliana* seedlings were grown for 5 days on half-strength Murashige and Skoog (MS) medium containing 1% agar (BD). *L. japonicus* seedlings were grown for 6 days on half-strength Hoagland medium containing 2.5μM PO_4_^3-^ and 0.4% Gelrite (www.duchefa-biochemie.com), or on half-strength MS containing 0.8% agar (only for experiment in Fig. 6D). Long-day conditions with 16h/8h light/dark cycles were used to test restoration of *A. thaliana* hypocotyl growth suppression by cross-species complementation (Fig. 2A). For Karrikin, *rac*-GR24, GR24^5DS^ and GR24^*ent*-5DS^ treatments the medium was supplied with KAR_1_ (www.olchemim.cz), KAR_2_ (www.olchemim.cz), *rac*-GR24 (www.chiralix.com) GR24^5DS^ and GR24^*ent*-5DS^ (www.strigolab.eu) or equal amounts of the corresponding solvent as a control. Karrikins were solubilized in 75% methanol and *rac*-GR24 and the GR24 stereoisomers in 100% acetone, at 10mM stock solution. Short-day conditions at 8h/16h light/dark cycles were used to test hormone responsiveness of *A. thaliana* as well as *L. japonicus* hypocotyls. After high-resolution scanning, the hypocotyl length was measured with Fiji (http://fiji.sc/).

### Root system architecture assay

*L. japonicus* germinated seeds were transferred onto new plates containing KAR_1_ (www.olchemim.cz), KAR_2_ (www.olchemim.cz), *rac*-GR24 (www.chiralix.com) or the corresponding solvent. Karrikins were solubilized in 75% methanol and *rac*-GR24 in 100% acetone, at 10 mM stock solution. Plates were partially covered with black paper to keep the roots in the dark, and placed at 24°C with 16-h-light/8-h-dark cycles for 2 weeks. After high-resolution scanning, post-embryonic root number was counted and primary root length measured with Fiji (http://fiji.sc/).

### Treatment for analysis of transcript accumulation

Seedling roots were placed in 1/2 Hoagland solution with 2.5 μM PO_4_^3-^ containing 1 or 3 μM Karrikin1 (www.olchemim.cz for qPCR analysis, synthesized according to [75] for microarray analysis), Karrikin2 (www.olchemim.cz), *rac*-GR24 (www.chiralix.com) or equal amounts of the corresponding solvents for the time indicated in Figure legends and the roots were covered with black paper to keep them in the dark.

### Microarray analysis

Three biological replicates were performed for each treatment. Root tissues were harvested, rapidly blotted dry and shock frozen in liquid nitrogen. RNA was extracted using the Spectrum Plant Total RNA Kit (www.sigmaaldrich.com). RNA was quantified and evaluated for purity using a Nanodrop Spectrophotometer ND-100 (NanoDrop Technologies, Willington, DE) and Bioanalyzer 2100 (Agilent, Santa Clara, CA). For each sample, 500 ng of total RNA was used for the expression analysis of each sample using the Affymetrix GeneChip^®^ Lotus1a520343 (Affymetrix, Santa Clara, CA). Probe labeling, chip hybridization and scanning were performed according to the manufacturer’s instructions for IVT Express Labeling Kit (Affymetrix). The Microarray raw data was normalized with the Robust Multiarray Averaging method (RMA) [76] using the Bioconductor [77] package “Methods for Affymetrix Oligonucleotide Arrays” (affy version 1.48.0) [78]. Control and rhizobial probesets were removed before statistical analysis. Differential gene expression was analyzed with the Bioconductor package “Linear Models for Microarray Data” (LIMMA version 3.26.8) [79]. The package uses linear models for parameter estimation and an empirical Bayes method for differential gene expression assessment [80]. P-values were adjusted due to multiple comparisons with the Benjamini-Hochberg correction (implemented in the LIMMA package). Probesets were termed as significantly differentially expressed, if their adjusted p-value was smaller than or equal to 0.01 and the fold change for at least one contrast showed a difference of at least 50%. To identify the corresponding gene models, the probeset sequences were used in a BLAST search against *L. japonicus* version 2.5 CDS and version 3.0 cDNA sequences (http://www.kazusa.or.jp/lotus/). If, based on the bitscore, multiple identical hits were found, we took the top hit in version 2.5 CDS as gene corresponding to the probe. For version 3.0 cDNA search we used the best hit, that was not located on chromosome 0, if possible. For probesets known to target chloroplast genes (probeset ID starting with Lj_), we preferred the best hit located on the chloroplast chromosome, if possible. Probeset descriptions are based on the info file of the *L. japonicus* Microarray chip provided by the manufacturer (Affymetrix).

### qPCR analysis

Tissue harvest, RNA extraction, cDNA synthesis and qPCR were performed as described previously [71]. qPCR reactions were run on an iCycler (Bio-Rad, www.bio-rad.com) or on QuantStudio5 (Applied Biosystems, www.thermofisher.com). Expression values were calculated according to the ΔΔCt method [81]. Expression values were normalized to the expression level of the housekeeping gene *Ubiquitin*. For each condition three to four biological replicates were performed. Primers are indicated in Table S4.

### Statistics

Statistical analyses were performed using Rstudio (www.rstudio.com) after log transformation for qPCR analysis. F- and p-values for all figures are provided in S5 Table.

## Acknowledgments

We thank Andreas Keymer and Priya Pimprikar for cDNAs from root, flower, leaf and stem; and Verena Klingl for excellent technical support. We thank Martin Parniske (LMU Munich, Germany) for providing microarray chips and for setting up the *L. japonicus* mutant collection at the Sainsbury laboratory Norwich, UK; to Jens Stougaard and all scientists at LotusBase for the LORE1 insertion lines (University of Aarhus, Denmark); and to David Nelson (University of California Riverside, USA) for fruitful discussion. We thank Min Ni (University of Minnesota, USA) for seeds of the *A. thaliana htl-2* mutant. The study was supported by an Australian Research Council Future Fellowship (FT150100162) to M.T.W. and the Emmy Noether program of the DFG, grant GU1423/1-1 to C.G..

## Supplementary figure legends

**S1 Fig. MAX2-like underwent pseudogenization.**

(**A**) Schematic representation of the synthenic regions containing the *MAX2* and *MAX2-like* loci in *L. japonicus*. Coloured arrows and black lines show exons and introns respectively.

(**B**) Protein alignment of *Lj*MAX2, *Lj*MAX2-like and an artificial *Lj*MAX2-like with a deletion of the thymine at the position 453 in the coding sequence (*Lj*MAX2-like ΔT453). Position of the nucleotide deletion is indicated in the translated sequence by a red triangle. Amino-acid conservation between MAX2 and MAX2-like is indicated by a dark background.

**S2 Fig. Organ-specific accumulation of *D14, KAI2a, KAI2b* and *MAX2* transcripts.**

(**A-C**) Transcript accumulation in wild-type of *D14*, *KAI2a, KAI2b* and *MAX2* normalized to expression of *Ubiquitin*, in (**A**) leaf, stem, flower and root of plants grown in pots, and in (**B**) hypocotyl and roots of 1 wpg plants grown on Petri dishes in 8h light / 16h dark cycles, and in (**c**) roots of 2 wpg plants grown on Petri dishes in 16h light / 8h dark cycles (n = 3).

**S3 Fig. Subcellular localisation of *Lj*D14, *Lj*KAI2a, *Lj*KAI2b and *Lj*MAX2 in *Nicotiana benthamiana* leaves.**

(**A**) Subcellular localization of *Lj*D14, *Lj*KAI2a, *Lj*KAI2b and *Lj*MAX2 in *N. benthamiana* leaf epidermal cells. *Lj*D14, *Lj*KAI2a and *Lj*KAI2b are N-terminally fused with mOrange. *Lj*MAX2 is N-terminally fused with T-Sapphire. Scale bars: 25 μm. (**B**) Western blot of protein extracts from *N. benthamiana*, showing that the mOrange tag fused with *Lj*D14, *Lj*KAI2a and *Lj*KAI2b was not cleaved at detectable amounts.

**S4 Fig. SDS-PAGE of purified SUMO fusion proteins and DSF assay with GR24^5DS^.**

(**A**) 200 pmol (approx. 8 μg) of purified proteins were separated by 12% SDS-PAGE containing 2,2,2-trichlorethanol as a vizualization agent. Below each lane is the calculated protein size in kiloDaltons. S, protein size standards (Precision Plus Dual Color Standards, Bio-Rad #1610394) with corresponding sizes in kDa shown on the left. Optimal exposures of recombinant proteins and size standards were taken separately under UV transillumination and red epi-illumination, respectively. The two images were merged in post-processing, and the junction between them is indicated by a vertical line. (**B**) DSF curves of purified SUMO fusion proteins of wild-type *Lj*KAI2a and *Lj*KAI2b, and versions with swapped amino acids *Lj*KAI2a^W157,M160,L190^, *Lj*KAI2b^F158,L161,S191^, *Lj*KAI2a^W157^, *Lj*KAI2b^F158^, at the indicated concentrations of GR24^5DS^. The first derivative of the change of fluorescence was plotted against the temperature. Each curve is the arithmetic mean of four technical replicates. Peaks indicate the protein melting temperature. There is no ligand-induced thermal destabilisation consistent with no protein-ligand interaction.

**S5 Fig. Amino acid differences between the legume KAI2a and KAI2b clades.**

Protein sequence alignment of KAI2a and KAI2b homologs from the legumes *Lotus japonicus, Pisum sativum, Medicago truncatula* and *Glycine max*, in comparison with Arabidopsis KAI2 and rice D14L. Residues conserved within the KAI2a and KAI2b clades but different between these clades are coloured in green and blue. Residues of the catalytic triad are coloured in red. A non-conserved tryptophan in *Lj*KAI2b located in the protein cavity is coloured in violet. Yellow triangles indicated amino acid residues located in the ligand-binding cavity of the proteins. Orange triangles indicate the three amino acids responsible for differences in GR24^*ent*-5DS^-binding between *Lj*KAI2a and *Lj*KAI2b.

**S6 Fig. Intrinsic tryptophan fluorescence assay confirms inability of *Lj*KAI2b to interact with GR24^*ent*-5DS^**

Intrinsic tryptophane fluorescence of wild-type *Lj*KAI2a and *Lj*KAI2b, and protein versions with swapped amino acids *Lj*KAI2a^M160,L190^, *Lj*KAI2b^L161,S191^, *Lj*KAI2a^W157,M160,L190^, *Lj*KAI2b^F158,L161,S191^, *Lj*KAI2a^W157^, *Lj*KAI2b^F158^ measured with (**A**) fixed wavelength filters (excitation 295/10 nm; longpass dichroic 325 nm; emission 360/20 nm) and (**B**) with a linear variable filter monochromator for emission wavelength scans (excitation 295/10 nm, emission 334-400 nm, step width 2 nm, emission bandwidth 8 nm) at the indicated GR24^*ent*-5DS^ concentrations.

**S7 Fig. The F157 to W replacement occured multiple times in angiosperm KAI2 proteins.**

Phylogenetic tree of KAI2 proteins rooted with *A. thaliana* DLK2. The KAI2a and KAI2b clades in legumes are highlighted by red and blue branches. Monophyletic groups corresponding to a same order or clade are highlighted by colored rectangular boxes. Amino-acids at the positions corresponding to AtKAI2 157, 160 and 190 are indicated with single-letter code. A black background indicates the presence of the most common residues in KAI2 proteins: F157, L160 and A190. A blue background indicates residues M160 and L190, conserved in legume KAI2b. A red background indicates S190, conserved in legume KAI2a. A green background indicates a W at position 157. A brown background indicates a different residue.

**S8 Fig. Transcript accumulation in the *L. japonicus* KAR and SL receptor mutants.**

(**A**) qRT-PCR based transcript accumulation of *LjKAI2a* and *LjKAI2b*, in roots of wild type and *kai2a-1, kai2b-1, kai2b-3, kai2a-1 kai2b-1* and *max2-4* as well as *LjMAX2* and *LjD14* in *max2-4* and *d14-1*, respectively (n=4). Expression values were normalized to those of the housekeeping gene *Ubiquitin*. (**B**) *LjKAI2b* transcript accumulation in wild-type, *kai2b-1* (stop codon) and *kai2b-3* (LORE1 insertion) mutants by semi-quantitative RT-PCR using primer pairs located 5’ and 3’ of the mutations, as well as flanking (ML) the mutations. Transcript accumulation of the housekeeping gene Ubiquitin is also shown.

**S9 Fig. Characterisation of the *kai2a-1* allele.**

(**A**) Schematic representation of mis-splicing caused by the LORE1 insertion in the *kai2a-1* mutant. (**B**) cDNA alignment showing the absence of nucleotides 369 to 383 in the *kai2a-1* transcript, causing a deletion of amino acids 124 to 128 (orange). (**C**) Protein model of *Lj*KAI2a based on the *At*KAI2-KAR_1_ complex 4JYM [5] showing KAR_1_ in green, residues of the catalytic triad in red and the amino acids missing in a hypothetical *Lj*KAI2a-1 protein in orange. (**D**) Hypocotyl elongation at 6 dpg in Arabidopsis *kai2-2* mutants transgenically complemented with genomic and the cDNA of wild-type *LjKAI2a* and *Ljkai2a-1* driven by the *AtKAI2* promoter (n = 75-106). Plants were grown in 8h light / 16h dark cycles. Letters indicate different statistical groups (ANOVA, post-hoc Tukey test).

**S10 Fig. *Lotus japonicus* hypocotyls respond to KAR_1_ and KAR_2_ in a *LjKAI2a*- and *LjMAX2*-dependent manner.**

(**A**) Hypocotyls and (**B**) hypocotyl length of *L. japonicus* seedling at 1 wpg after treatment with solvent (M) or three different concentrations of KAR_1_, KAR_2_ or *rac*-GR24 (GR24) (n= 95-105). Letters indicate different statistical groups (ANOVA, post-hoc Tukey test). (**C**) Hypocotyl length of the indicated genotypes at 1 wpg after treatment with solvent (Mock), 1 μM KAR_1_ or 1 μM KAR_2_ (n = 73-107). (**D**) Hypocotyl length of wild-type and *max2-4* seedlings 1 wpg after treatment with solvent (Mock), 1 μM KAR_1_, 1 μM KAR_2_ (n = 66-96). (**E**) RT-qPCR-based expression of *DLK2* in hypocotyls at 1 wpg after 2 hours treatment with solvent (Mock), 1 μM KAR_1_, 1 μM KAR_2_, or 1 μM *rac*-GR24 (GR24) (n = 3). Expression values were normalized to those of the housekeeping gene *Ubiquitin*. (**A-E**) Seedlings were grown in 8h light / 16h dark cycles. (**C-E**) Asterisks indicate significant differences of the compounds versus mock treatment (ANOVA, post-hoc Dunnett test, N.S.>0.05, *≤0.05, **≤0.01, ***≤0.001).

**S11 Fig. Small overlap between transcriptional responses of *Lotus japonicus* roots to KAR_1_ and *rac*-GR24.**

Number of differentially expressed genes (DEGs, adjusted p-value < 0.01) as assessed by microarray analysis. Left panel: DEGs responding to 1 μM KAR_1_ after 1h, 2h and 6h incubation. Middle panel: DE genes responding to 1 μM *rac*-GR24 1h, 2, 6h incubation. Right panel: comparison of DE genes responding to 2 h treatment with KAR_1_ and *rac*-GR24.

**S12 Fig. KAR perception mutants are less responsive to KAR_1_ treatment.**(**A-C**) Post-embryonic-root (PER) density of *L. japonicus* plants, 2 wpg after treatment with solvent (Mock) or 1 μM KAR_1_, of wild-type, (**A**) *kai2a-1, kai2b-1* and *kai2a-1 kai2b-1* (n= 3250); (**B**) *max2-4* (n= 34-43); (**c**) *kai2a-1, kai2b-3* and *kai2a-1 kai2b-1* (n= 37-72). (**A-C**) Asterisks indicate significant differences versus mock treatment (Welch t.test, *≤0.05, **≤0.01, ***≤0.001).

**S13 Fig. KAR_1_ response in roots requires *LjKAI2a* or *LjKAI2b* and *LjMAX2*.**

Primary-root length (PRL) and post-embryonic-root (PER) number of *L. japonicus* plants, 2 wpg after treatment with solvent (Mock) or 3 μM KAR_1_ (n=34-72) displayed in Fig 9A. Asterisks indicate significant differences versus mock treatment (Welch t.test, *≤0.05, **≤0.01, ***≤0.001).

## Notes

### Competing Interest Statement

The authors have declared no competing interest.

