## Supplemental Figures and Tables for "*Lotus japonicus* karrikin receptors display divergent ligand-binding specificities and organ-dependent redundancy"

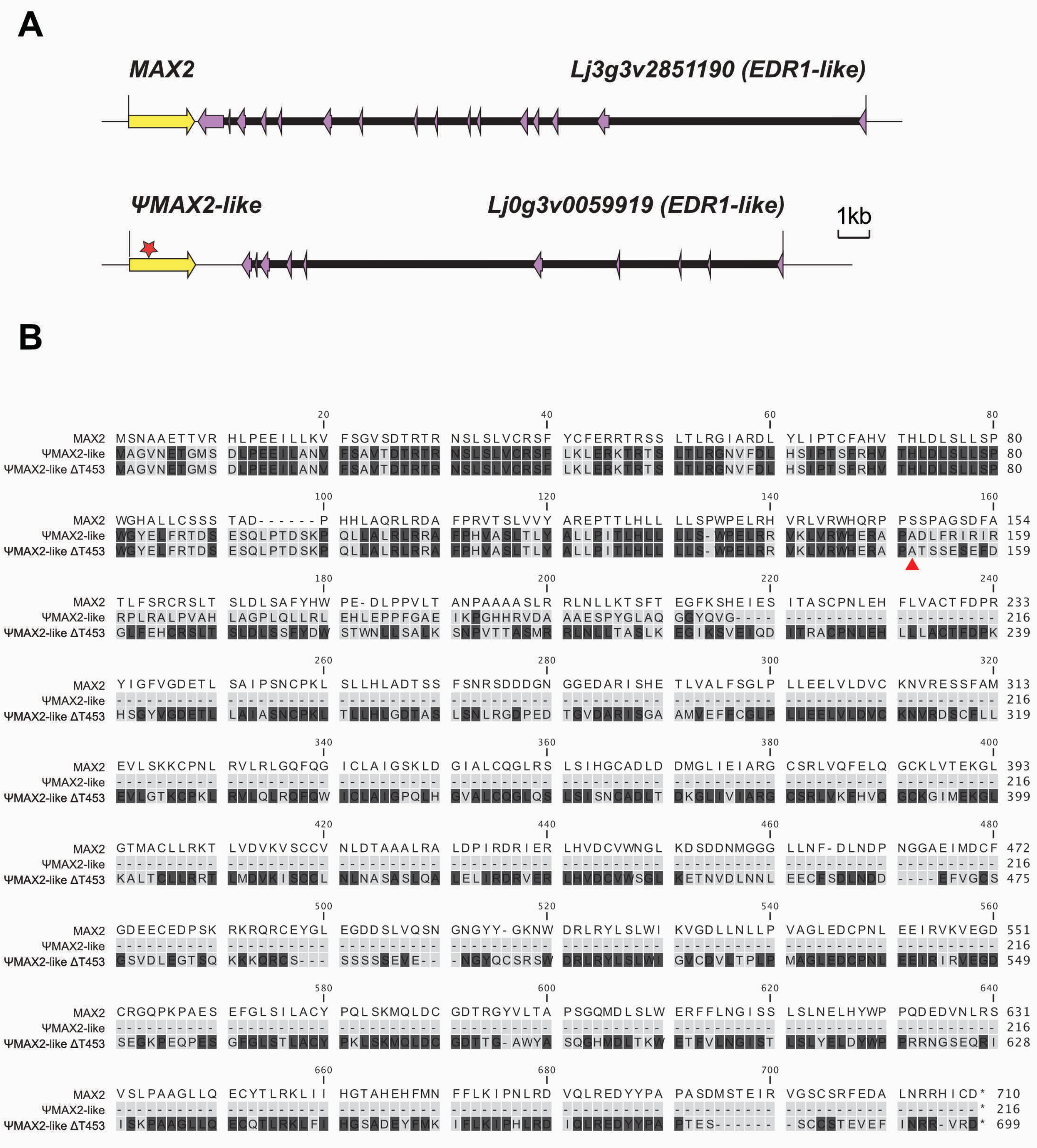


**S1 Fig. MAX2-like underwent pseudogenization.**
(**A**) Schematic representation of the synthenic regions containing the *MAX2* and *MAX2-like* loci in *L. japonicus*. Coloured arrows and black lines show exons and introns respectively. (**B**) Protein alignment of *Lj*MAX2, *Lj*MAX2-like and an artificial *Lj*MAX2-like with a deletion of the thymine at the position 453 in the coding sequence (*Lj*MAX2-like ΔT453). Position of the nucleotide deletion is indicated in the translated sequence by a red triangle. Amino-acid conservation between MAX2 and MAX2-like is indicated by a dark background.

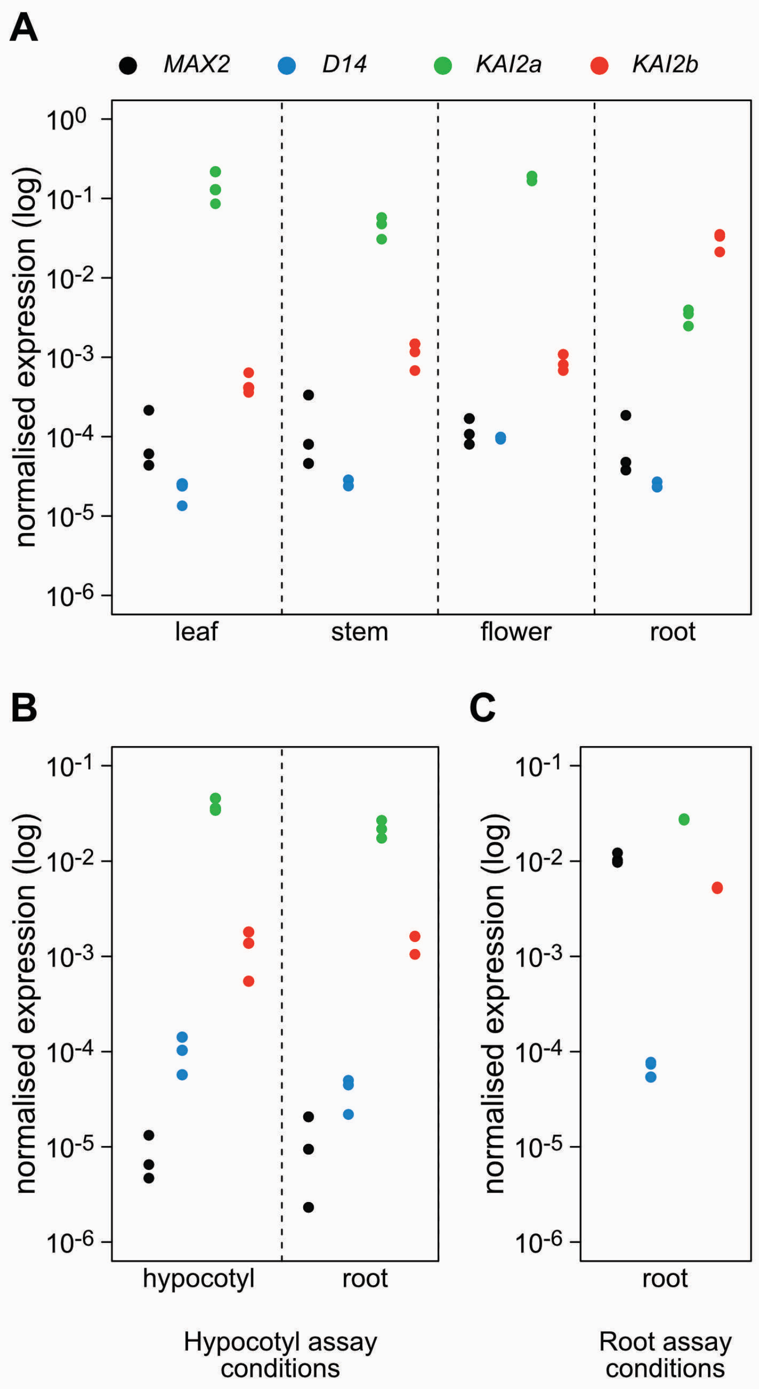


**S2 Fig. Organ-specific accumulation of *D14*, *KAI2a*, *KAI2b* and *MAX2* transcripts.**
(**A-C**) Transcript accumulation in wild-type of *D14*, *KAI2a*, *KAI2b* and *MAX2* normalized to expression of *Ubiquitin*, in (**A**) leaf, stem, flower and root of plants grown in pots, and in (**B**) hypocotyl and roots of 1 wpg plants grown on Petri dishes in 8h light / 16h dark cycles, and in (**c**) roots of 2 wpg plants grown on Petri dishes in 16h light / 8h dark cycles (n = 3).

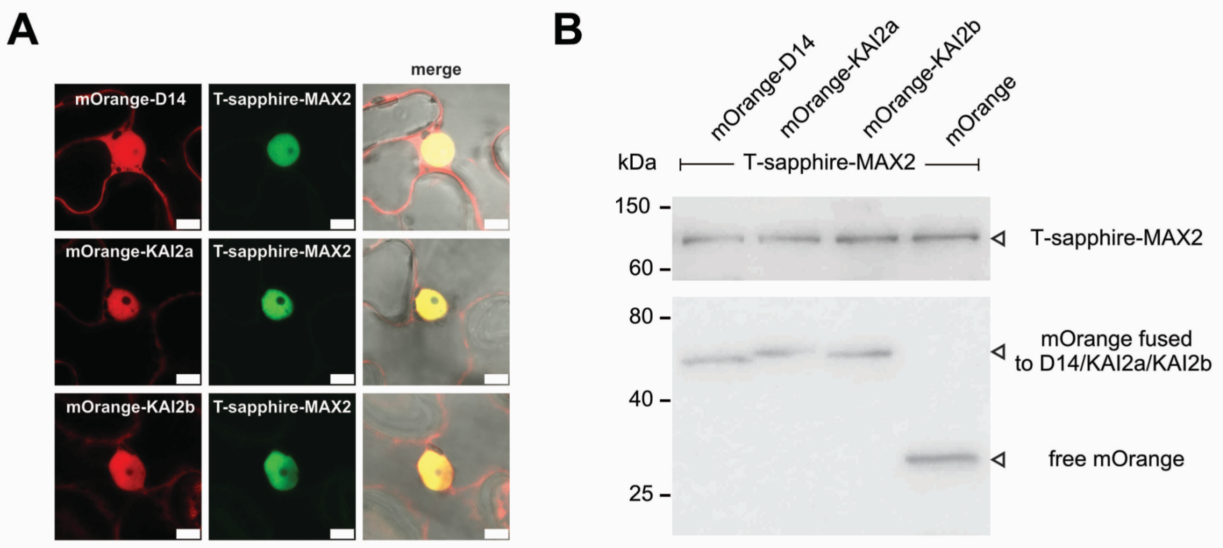


**S3 Fig. Subcellular localisation of *Lj*D14, *Lj*KAI2a, *Lj*KAI2b and *Lj*MAX2 in *Nicotiana benthamiana* leaves.**
(**A**) Subcellular localization of *Lj*D14, *Lj*KAI2a, *Lj*KAI2b and *Lj*MAX2 in *N. benthamiana* leaf epidermal cells. *Lj*D14, *Lj*KAI2a and *Lj*KAI2b are N-terminally fused with mOrange. *Lj*MAX2 is N-terminally fused with T-Sapphire. Scale bars: 25 µm. (**B**) Western blot of protein extracts from *N. benthamiana*, showing that the mOrange tag fused with *Lj*D14, *Lj*KAI2a and *Lj*KAI2b was not cleaved at detectable amounts.

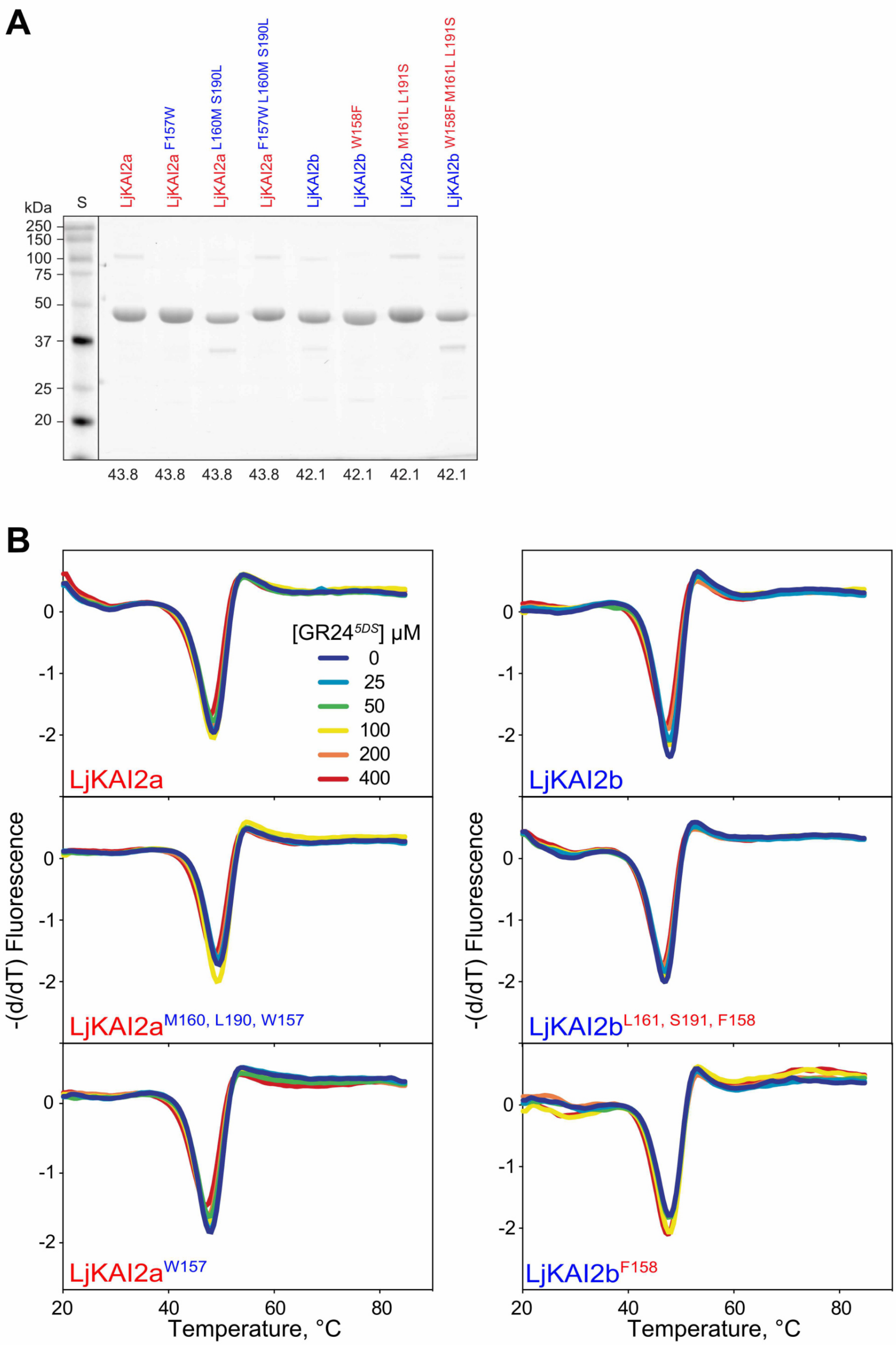


**S4 Fig. SDS-PAGE of purified SUMO fusion proteins and DSF assay with GR24$^{5DS}$.**

(**A**) 200 pmol (approx. 8 µg) of purified proteins were separated by 12% SDS-PAGE containing 2,2,2-trichlorethanol as a vizualization agent. Below each lane is the calculated protein size in kiloDaltons. S, protein size standards (Precision Plus Dual Color Standards, Bio-Rad #1610394) with corresponding sizes in kDa shown on the left. Optimal exposures of recombinant proteins and size standards were taken separately under UV transillumination and red epi-illumination, respectively. The two images were merged in post-processing, and the junction between them is indicated by a vertical line. (**B**) DSF curves of purified SUMO fusion proteins of wild-type *Lj*KAI2a and *Lj*KAI2b, and versions with swapped amino acids *Lj*KAI2a$^{W157,M160,L190}$, *Lj*KAI2b$^{F158,L161,S191}$, *Lj*KAI2a$^{W157}$, *Lj*KAI2b$^{F158}$, at the indicated concentrations of GR24$^{5DS}$. The first derivative of the change of fluorescence was plotted against the temperature. Each curve is the arithmetic mean of four technical replicates. Peaks indicate the protein melting temperature. There is no ligand-induced thermal destabilisation consistent with no protein-ligand interaction.

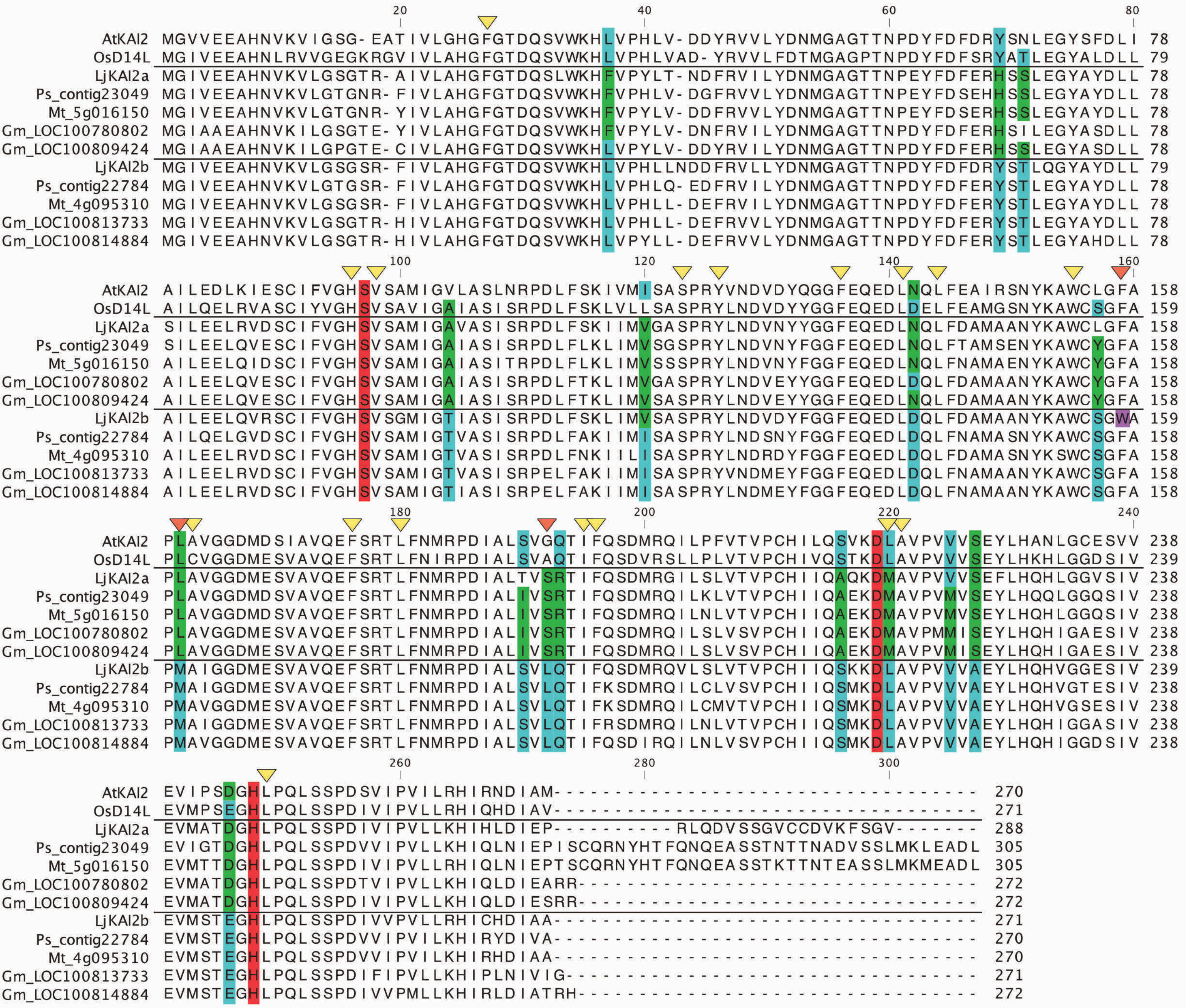


**S5 Fig. Amino acid differences between the legume KAI2a and KAI2b clades.**

Protein sequence alignment of KAI2a and KAI2b homologs from the legumes *Lotus japonicus, Pisum sativum, Medicago truncatula* and *Glycine max*, in comparison with Arabidopsis KAI2 and rice D14L. Residues conserved within the KAI2a and KAI2b clades but different between these clades are coloured in green and blue. Residues of the catalytic triad are coloured in red. A non-conserved tryptophan in *Lj*KAI2b located in the protein cavity is coloured in violet. Yellow triangles indicated amino acid residues located in the ligand-binding cavity of the proteins. Orange triangles indicate the three amino acids responsible for differences in GR24$^{ent\text{-}5DS}$-binding between *Lj*KAI2a and *Lj*KAI2b.

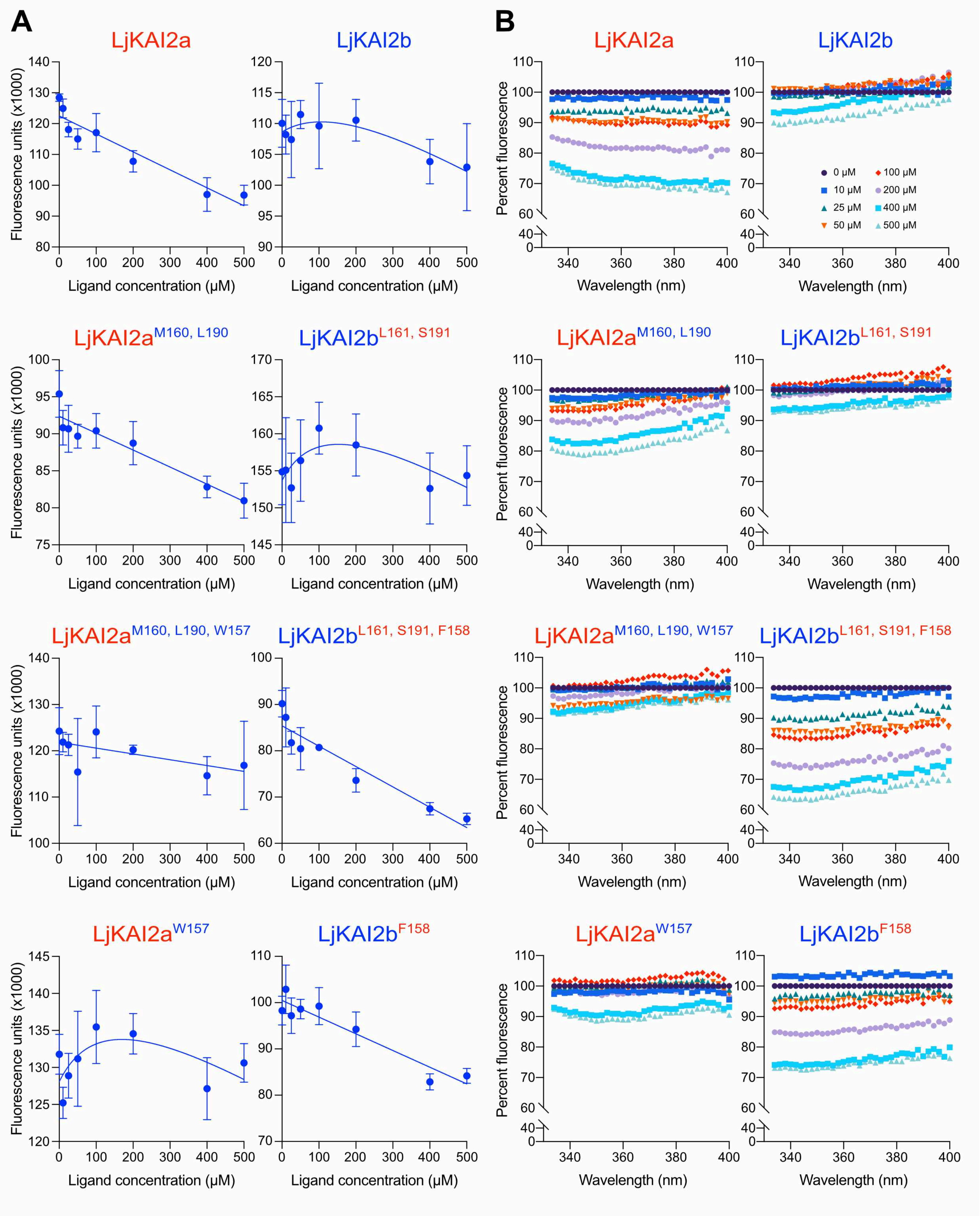


**S6 Fig. Intrinsic tryptophan fluorescence assay confirms inability of *Lj*KAI2b to interact with GR24*ent*-5DS**

Intrinsic tryptophane fluorescence of wild-type *Lj*KAI2a and *Lj*KAI2b, and protein versions with swapped amino acids *Lj*KAI2a$^{M160,L190}$, *Lj*KAI2b$^{L161,S191}$, *Lj*KAI2a$^{M160,L190,W157}$, *Lj*KAI2b$^{L161,S191,F158}$, *Lj*KAI2a$^{W157}$, *Lj*KAI2b$^{F158}$ measured with (**A**) fixed wavelength filters (excitation 295/10 nm; longpass dichroic 325 nm; emission 360/20 nm) and (**B**) with a linear variable filter monochromator for emission wavelength scans (excitation 295/10 nm, emission 334-400 nm, step width 2 nm, emission bandwidth 8 nm) at the indicated GR24$^{ent\text{-}5DS}$ concentrations.

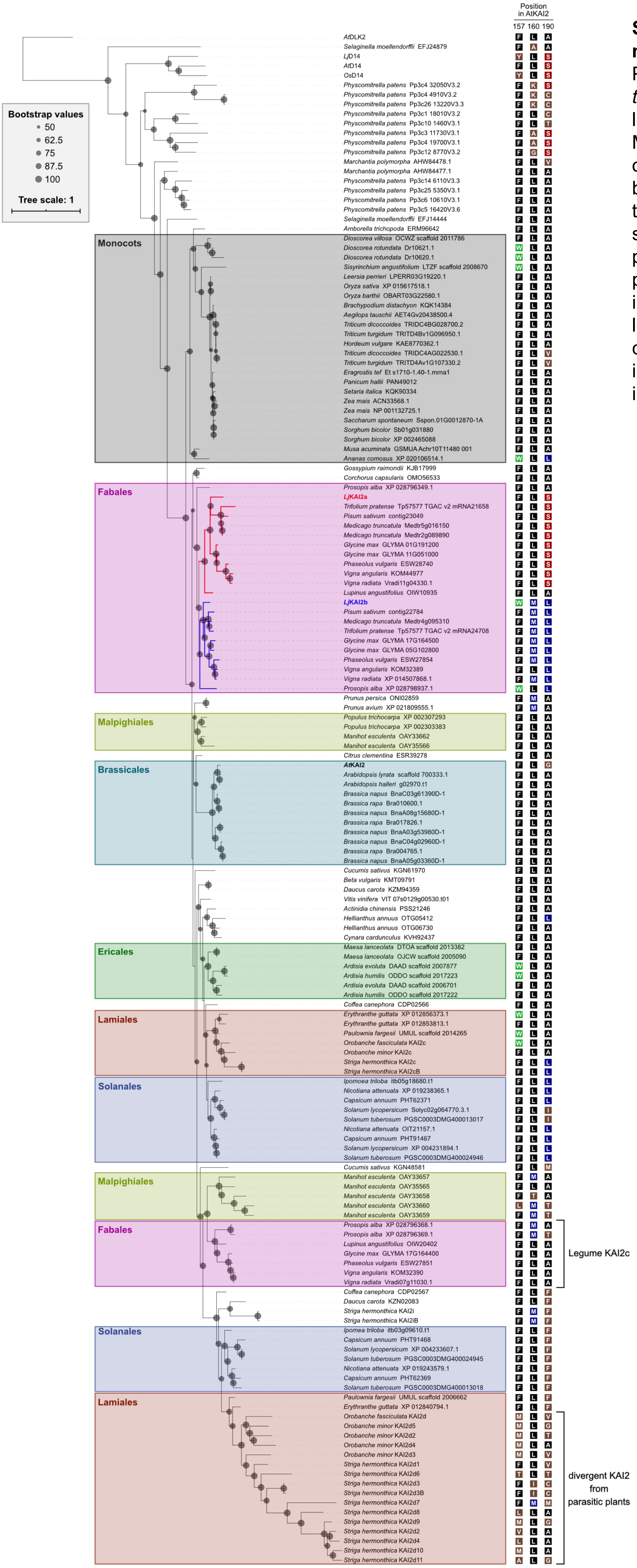


**S7 Fig. The F157 to W replacement occured multiple times in angiosperm KAI2 proteins.** Phylogenetic tree of KAI2 proteins rooted with *A. thaliana* DLK2. The KAI2a and KAI2b clades in legumes are highlighted by red and blue branches. Monophyletic groups corresponding to a same order or clade are highlighted by colored rectangular boxes. Amino-acids at the positions corresponding to AtKAI2 157, 160 and 190 are indicated with single-letter code. A black background indicates the presence of the most common residues in KAI2 proteins: F157, L160 and A190. A blue background indicates residues M160 and L190, conserved in legume KAI2b. A red background indicates S190, conserved in legume KAI2a. A green background indicates a W at position 157. A brown background indicates a different residue.

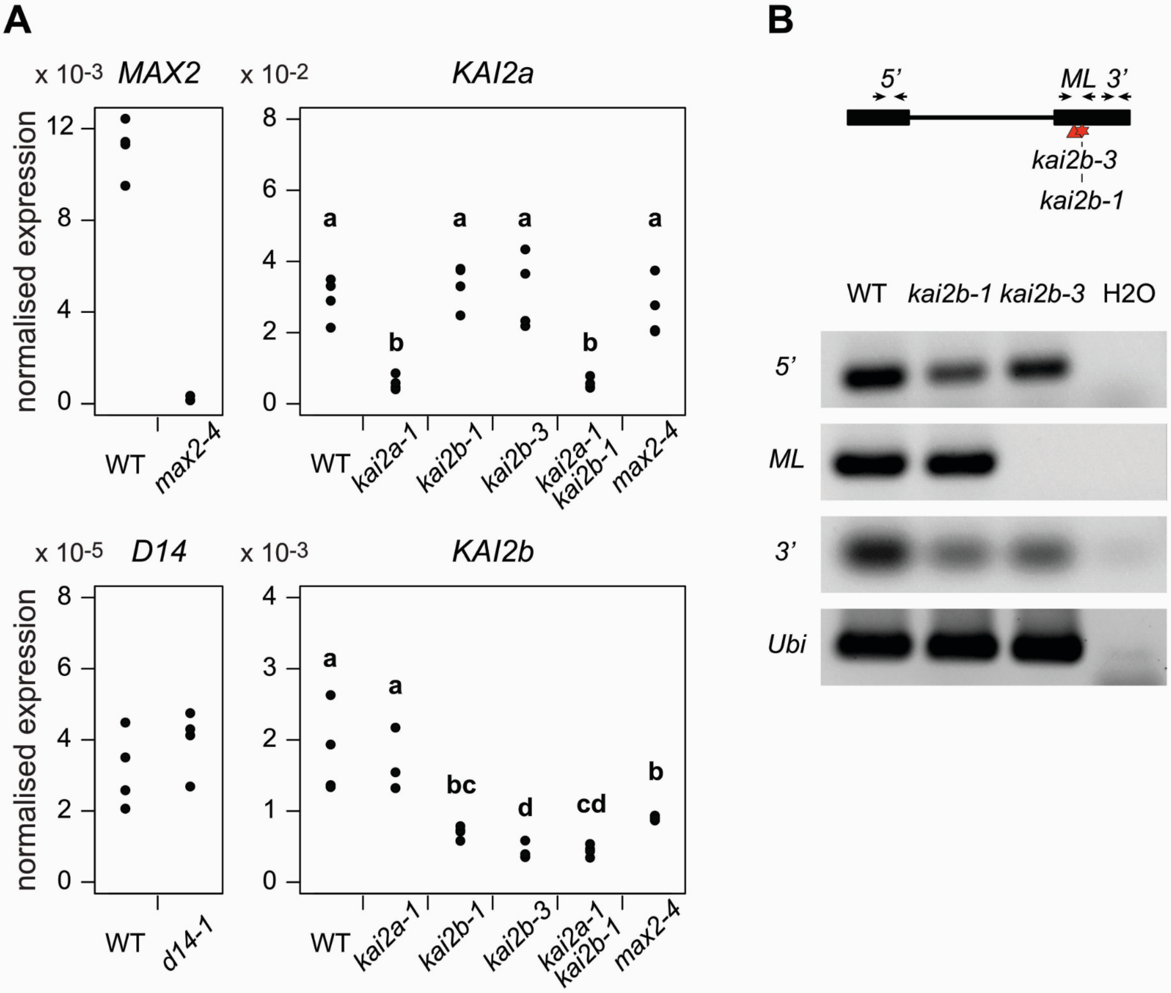


**S8 Fig. Transcript accumulation in the *L. japonicus* KAR and SL receptor mutants.**
(**A**) qRT-PCR based transcript accumulation of *LjKAI2a* and *LjKAI2b*, in roots of wild type and *kai2a-1*, *kai2b-1*, *kai2b-3*, *kai2a-1 kai2b-1* and *max2-4* as well as *LjMAX2* and *LjD14* in *max2-4* and *d14-1*, respectively (n=4). Expression values were normalized to those of the housekeeping gene *Ubiquitin*. (**B**) *LjKAI2*b transcript accumulation in wild-type, *kai2b-1* (stop codon) and *kai2b-3* (LORE1 insertion) mutants by semi-quantitative RT-PCR using primer pairs located 5' and 3' of the mutations, as well as flanking (ML) the mutations. Transcript accumulation of the housekeeping gene Ubiquitin is also shown.

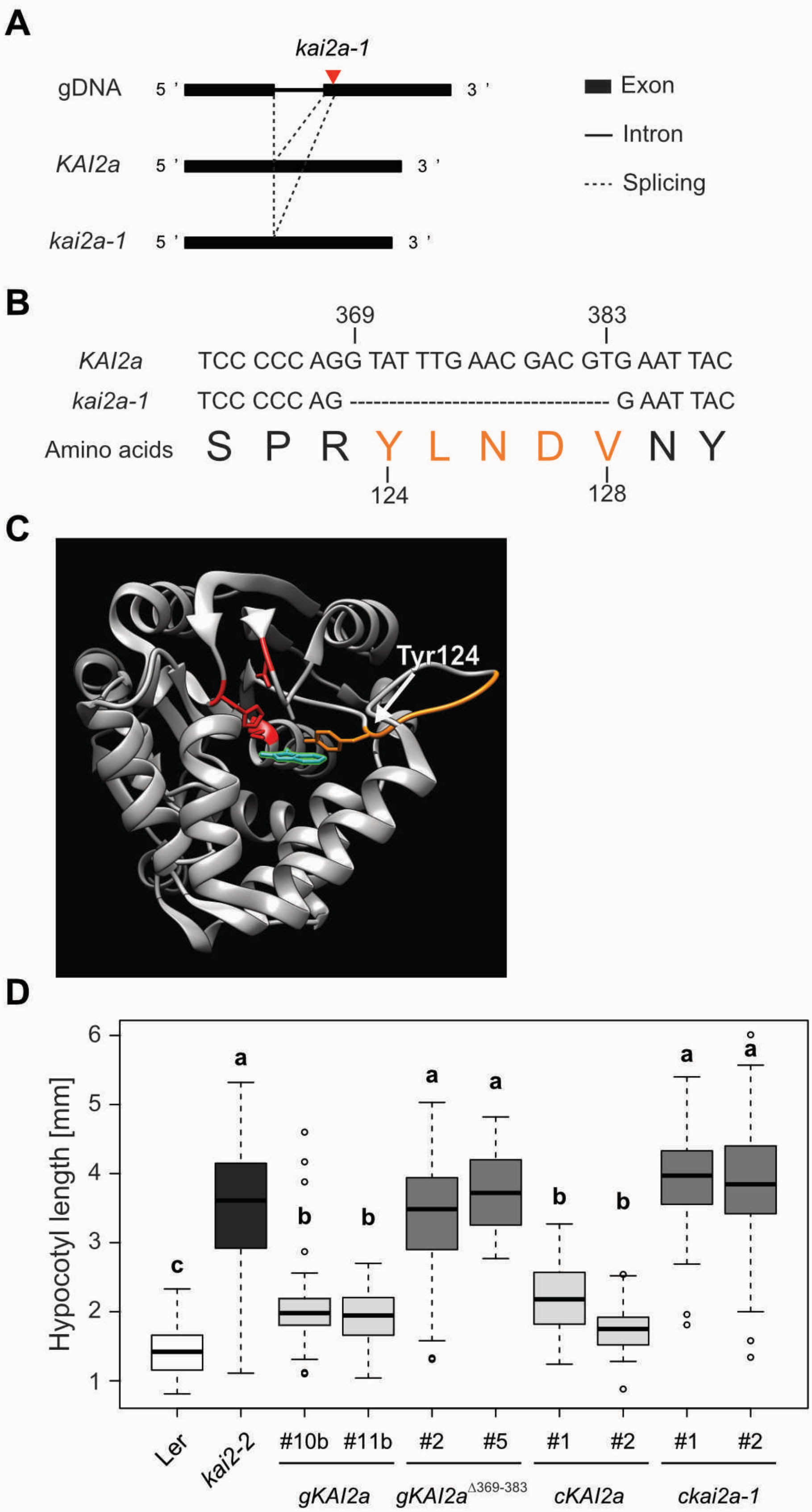


**S9 Fig. Characterisation of the *kai2a-1* allele.**

(**A**) Schematic representation of mis-splicing caused by the LORE1 insertion in the *kai2a-1* mutant. (**B**) cDNA alignment showing the absence of nucleotides 369 to 383 in the *kai2a-1* transcript, causing a deletion of amino acids 124 to 128 (orange). (**C**) Protein model of *Lj*KAI2a based on the *At*KAI2-KAR$_1$ complex 4JYM [5] showing KAR$_1$ in green, residues of the catalytic triad in red and the amino acids missing in a hypothetical *Lj*KAI2a-1 protein in orange. (**D**) Hypocotyl elongation at 6 dpg in Arabidopsis *kai2-2* mutants transgenically complemented with genomic and the cDNA of wild-type *LjKAI2a* and *Ljkai2a-1* driven by the *AtKAI2* promoter (n = 75-106). Plants were grown in 8h light / 16h dark cycles. Letters indicate different statistical groups (ANOVA, post-hoc Tukey test).

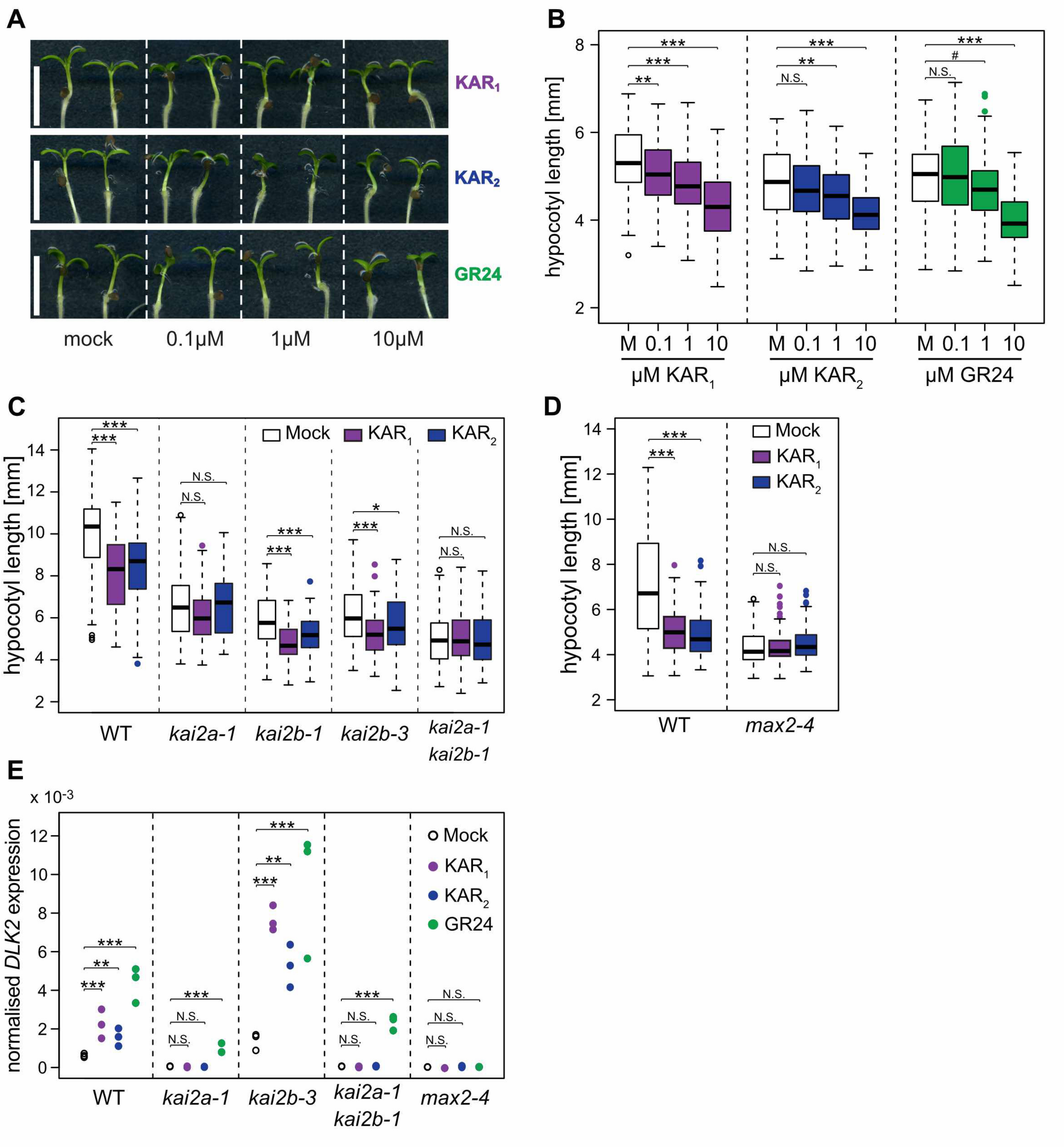


**S10 Fig. *Lotus japonicus* hypocotyls respond to $KAR_1$ and $KAR_2$ in a *LjKAI2a*- and *LjMAX2*-dependent manner.**
(**A**) Hypocotyls and (**B**) hypocotyl length of *L. japonicus* seedling at 1 wpg after treatment with solvent (M) or three different concentrations of $KAR_1$, $KAR_2$ or *rac*-GR24 (GR24) (n= 95-105). Letters indicate different statistical groups (ANOVA, post-hoc Tukey test). (**C**) Hypocotyl length of the indicated genotypes at 1 wpg after treatment with solvent (Mock), 1 µM $KAR_1$ or 1 µM $KAR_2$ (n = 73-107). (**D**) Hypocotyl length of wild-type and *max2-4* seedlings 1 wpg after treatment with solvent (Mock), 1 µM $KAR_1$, 1 µM $KAR_2$ (n = 66-96). (**E**) RT-qPCR-based expression of *DLK2* in hypocotyls at 1 wpg after 2 hours treatment with solvent (Mock), 1 µM $KAR_1$, 1 µM $KAR_2$, or 1 µM *rac*-GR24 (GR24) (n = 3). Expression values were normalized to those of the housekeeping gene *Ubiquitin*. (**A-E**) Seedlings were grown in 8h light / 16h dark cycles. (**C-E**) Asterisks indicate significant differences of the compounds versus mock treatment (ANOVA, post-hoc Dunnett test, N.S.>0.05, *≤0.05, **≤0.01, ***≤0.001).

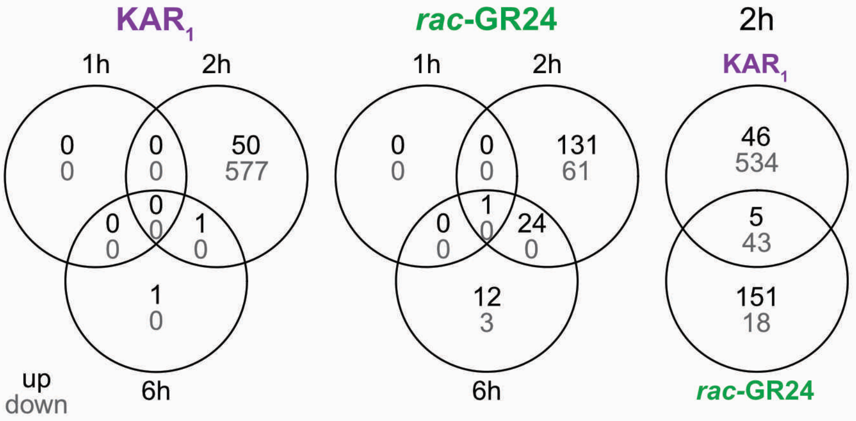


**S11 Fig. Small overlap between transcriptional responses of *Lotus japonicus* roots to KAR$_1$ and *rac*-GR24.**
Number of differentially expressed genes (DEGs, adjusted p-value < 0.01) as assessed by microarray analysis. Left panel: DEGs responding to 1 µM KAR$_1$ after 1h, 2h and 6h incubation. Middle panel: DE genes responding to 1 µM *rac*-GR24 1h, 2, 6h incubation. Right panel: comparison of DE genes responding to 2 h treatment with KAR$_1$ and *rac*-GR24.

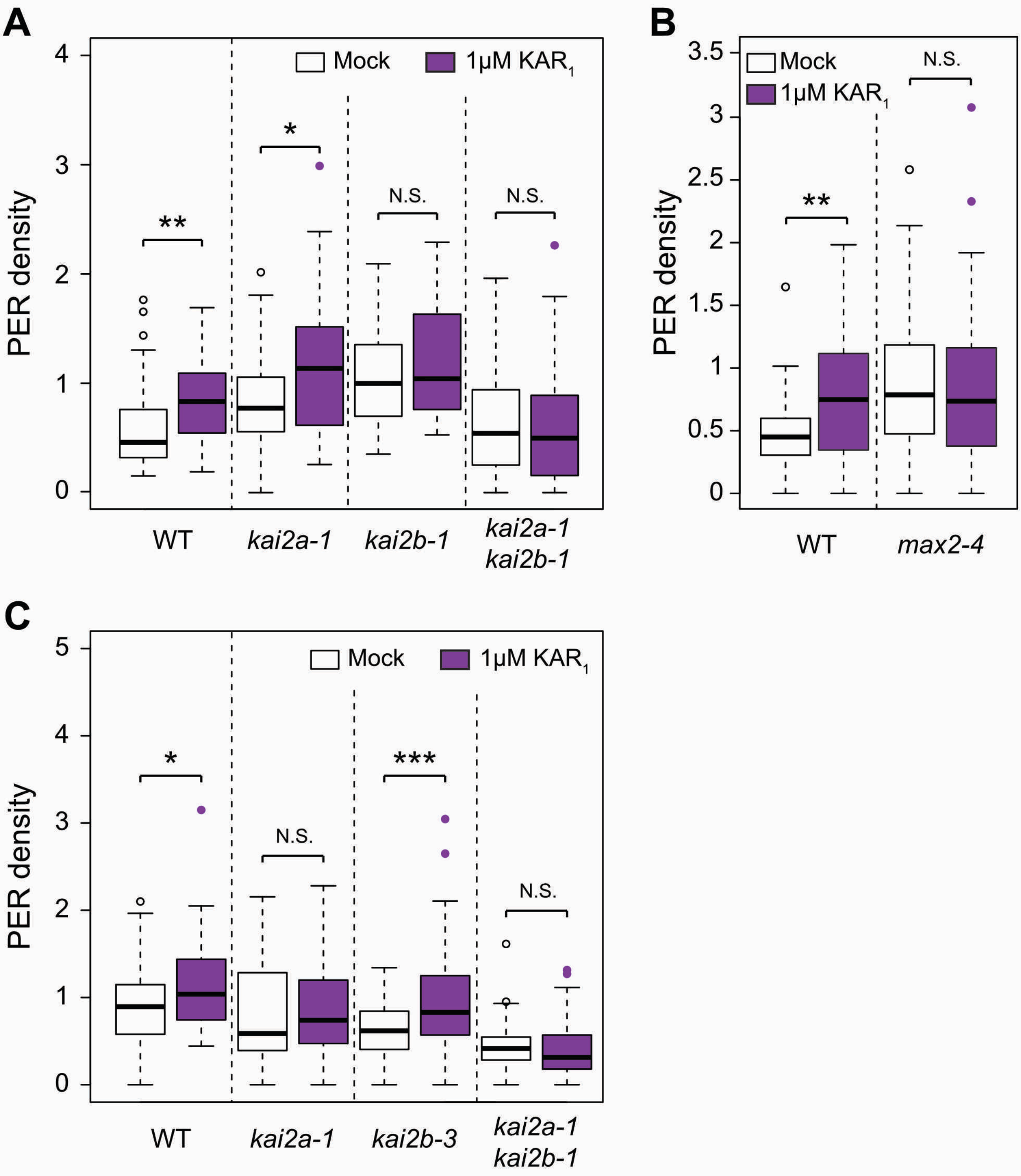


**S12 Fig. KAR perception mutants are less responsive to $KAR_1$ treatment.**
(**A-C**) Post-embryonic-root (PER) density of *L. japonicus* plants, 2 wpg after treatment with solvent (Mock) or 1 µM $KAR_1$, of wild-type, (**A**) *kai2a-1*, *kai2b-1* and *kai2a-1 kai2b-1* (n= 32-50); (**B**) *max2-4* (n= 34-43); (**c**) *kai2a-1*, *kai2b-3* and *kai2a-1 kai2b-1* (n= 37-72). (**A-C**) Asterisks indicate significant differences versus mock treatment (Welch t.test, *≤0.05, **≤0.01, ***≤0.001).

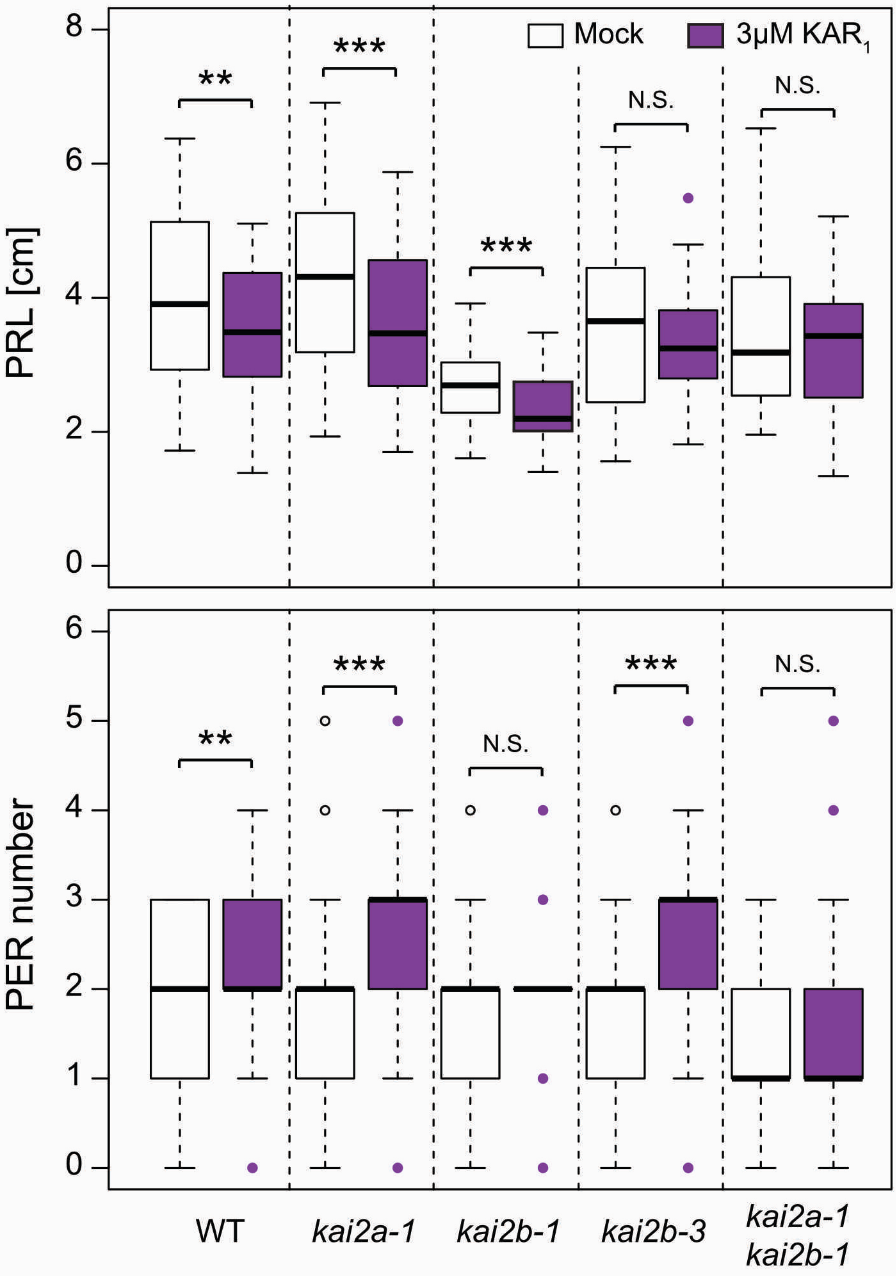


**S13 Fig. $KAR_1$ response in roots requires *LjKAI2a* or *LjKAI2b* and *LjMAX2*.**
Primary-root length (PRL) and post-embryonic-root (PER) number of *L. japonicus* plants, 2 wpg after treatment with solvent (Mock) or 3 µM $KAR_1$ (n=34-72) displayed in Fig 9A. Asterisks indicate significant differences versus mock treatment (Welch t.test, *≤0.05, **≤0.01, ***≤0.001).

**S1 Table.** *L. japonicus* mutants used in this study and information on seed production.

| allele | type | reference | position from ATG | comments |
|---|---|---|---|---|
| *Ljd14-1* | EMS | SL4580 | C685T (Q > stop) | hardly produce flowers |
| *Ljkai2a-1* | LORE1 insertion | 30008990 | 387 | - |
| *Ljkai2b-1* | EMS | SL1281 | C640T (Q > stop) | - |
| *Ljkai2b-2* | EMS | SL2723 | G462A (W > stop) | produced no seeds |
| *Ljkai2b-3* | LORE1 insertion | 30034333 | 535 | - |
| *Ljmax2-1* | LORE1 insertion | 30031159 | 83 | hardly produce flowers |
| *Ljmax2-2* | LORE1 insertion | P0860_3 | 504 | hardly produce flowers |
| *Ljmax2-3* | LORE1 insertion | 30019601 | 1132 | produce few flowers |
| *Ljmax2-4* | LORE1 insertion | 30049531 | 1230 | produce few flowers |

**S3 Table.** Primers

**Primers used for LORE1 insertion mutant genotyping.** Forward primer was used to amplify specifically LORE1 insertion with a specific P2 primer (CCATGGCGGTTCCGTGAATCTTAGG).

| mutant | Forward | | Reverse | |
|---|---|---|---|---|
| *Ljkai2a-1* | Sc403 | TATGGTCTCTCACGCTGTTTCCGCC ATGATCG | Sc283 | TCCACAATAGACACGCCACC |
| *Ljkai2b-3* | Sc285 | CCTCCGTTGACATGACCTCC | Sc17 | TTGAAGACTACCCCTTAAACA AGGGGTTTGAG |
| *Ljmax2-1* | Sc130 | ATGAAGACTTTACGGGTCTCACACC ATGAGTAACGCTGCTGAAAC | CG416 | CAGTAGAAGCTCCGGCAAAC |
| *Ljmax2-2* | CG383 | TTGGGGAGGGGTTTAATAGG | CG424 | CGATTTCGTGAGACTTGAAGC |
| *Ljmax2-3* | Sc163 | TCACCTCGCTGGATCTCTC | Sc131 | TTGAAGACTACCACCTCCCAT GTTGTCATC |
| *Ljmax2-4* | Sc131 | TTGAAGACTACCACCTCCCATGTTG TCATC | Sc163 | TCACCTCGCTGGATCTCTC |

**Primers used for EMS mutant genotyping.** dCAPS strategy was used to genotype EMS mutants.

| mutant | Forward | | Reverse | | Site |
|---|---|---|---|---|---|
| *Ljd14-1* | Sc429 | GCCGGCGGCGGCCGCGAGGT ACCTG | Sc242 | TTTCGTCTCACCTTGTGTGCCC CCGCCAGTGC | PstI (Cut WT) |
| *Ljkai2b-1* | Sc431 | GGTAACTGTGCCATGTCACAG TATA | Sc285 | CCTCCGTTGACATGACCTCC | AccI (Cut WT) |

**Primers used for cloning.**

| Use | Primers | |
|---|---|---|
| cloning promoter *AtD14* in LI | Sc224 | TTTCGTCTCAGCGGGTCTACACATTCATCAATCTCGC |
| | Sc225 | TTTCGTCTCACAGATTTTTTATGTGTTTGGGTTTGAG |
| cloning promoter *AtKAI2* fragment 1 in LI | Sc232 | TTTCGTCTCAGCGGGGCGATTCAGTGCCATGATT |
| | Sc233 | TTTCGTCTCACGATTCGTTCAGATTCTCGCT |
| cloning promoter *AtKAI2* fragment 2 in LI | Sc234 | TTTCGTCTCAATCGACTCGAATTTGATGGATCTTTC |
| | Sc235 | TTTCGTCTCACAGACTCTCTAAAGAAGATTCTTC |
| cloning genomic *AtD14* in LI | Sc236 | TTTCGTCTCACACCATGAGTCAACACAACATCTTAGAAG |
| | Sc237 | TTTCGTCTCACCTTTCACCGAGGAAGAGCTCGCC |
| cloning genomic *AtKAI2* in LI | Sc238 | TTTCGTCTCACACCATGGGTGTGGTAGAAGAAG |
| | Sc239 | TTTCGTCTCACCTTTCACATAGCAATGTCATTACGAATG |
| cloning genomic *LjD14* in LI | Sc240 | TTTCGTCTCACACCATGGCCACTTCAATCCTCGACG |
| | Sc241 | TTTCGTCTCACCTTTCAGTGTGCCCCCGCCAGTG |
| cloning genomic *LjKAI2a* and cDNA *Ljkai2a-1* in LI | Sc243 | TTTCGTCTCACACCATGGGGATAGTGGAGGAAGCTCAC |
| | Sc244 | TTTCGTCTCACCTTTTACACCCCACTAAATTTTACATCAC |

| | | |
|---|---|---|
| cloning genomic *LjKAI2b* in LI | Sc246 | TTTCGTCTCACACCATGGGGATAGTGGAAGAAGCTC |
| | Sc247 | TTTCGTCTCACCTTTCAAGCTGCAATATCATGGCAAATG |
| cloning genomic *Ljkai2a-1* in LI | Sc243 | TTTCGTCTCACACCATGGGGATAGTGGAGGAAGCTCAC |
| | ST97 | CAAATCCTTCCATAGTAATTTGCGGAAGAAAATCATC |
| | ST96 | TTGAAGACTATCTTCAGATATCTCATATAC |
| | Sc244 | TTTCGTCTCACCTTTTACACCCCACTAAATTTTACATCAC |
| cloning cDNA *LjKAI2a*$_{M160,L190,W157}$ fragment 1 in L0 | Sc505 | ATGAAGACTTCCATCGGAGCCCACCCTAAAC |
| | ST161 | ATGAAGACTTTACGTCGTCTCACACCATGGG |
| cloning cDNA *LjKAI2a*$_{M160,L190,W157}$ fragment 2 in L0 | ST163 | ATGAAGACTTATGGCGGTGGGTGGAGACATG |
| | ST164 | ATGAAGACTTCGCAAAACGGTTAGAGCAATATC |
| cloning cDNA *LjKAI2a*$_{M160,L190,W157}$ fragment 3 in L0 | ST165 | ATGAAGACTTTGCGGACCATTTTTCAGAGC |
| | Sc498 | ATGAAGACTACAGACGTCTCACCTTTTACACCCCACTAA ATTTTAC |
| cloning cDNA *LjKAI2b* $_{L161,S191,F158}$ fragment 1 in L0 | Sc506 | ATGAAGACTTCCAGCGGGGCAAAGCCTGAAC |
| | ST169 | ATGAAGACTTTACGTCGTCTCACACCATGGG |
| cloning cDNA *LjKAI2b* $_{L161,S191,F158}$ fragment 2 in L0 | ST171 | ATGAAGACTTCTGGCTATCGGAGGAGACATG |
| | ST172 | ATGAAGACTTTGCGATACGCTTAAGGCTATG |
| cloning cDNA *LjKAI2b* $_{L161,S191,F158}$ fragment 3 in L0 | ST173 | ATGAAGACTTCGCAGACAATTTTTCAAAGTG |
| | Sc503 | ATGAAGACTACAGACGTCTCACCTTTCAAGCTGCAATATC |
| cloning pSUMO *LjKAI2a*$_{M160,L190}$ | Sc604 | CGTGGTGTTTAGGGTTTGCTCCGATGGCGGTG |
| | Sc605 | CACCGCCATCGGAGCAAACCCTAAACACCACG |
| cloning pSUMO *LjKAI2b* $_{L161,S191}$ | Sc606 | CATGGTGTTCAGGCTGGGCCCCGCTGGCTATC |
| | Sc607 | GATAGCCAGCGGGGCCCAGCCTGAACACCATG |
| cloning pSUMO *LjKAI2a*$_{W157}$ | ST177 | GTGGTGTTTAGGGTGGGCTC |
| | ST178 | AGCGGAGCCCACCCTAAAC |
| cloning pSUMO *LjKAI2b* $_{F158}$ | ST179 | ATGGTGTTCAGGCTTTGC |
| | ST180 | ATAGCCATCGGGGCAAAG |

### Primers used for gene amplification by RT-qPCR.

| Use | Primers | |
|---|---|---|
| qPCR *Ubiquitin* | Ubi F | ATGCAGATCTTCGTCAAGACCTTG |
| | Ubi R | ACCTCCCCTCAGACGAAG |
| qPCR *LjMAX2* | Sc302 | GAATGTTACACCCTGAGGAAGC |
| | Sc303 | TCAGGTTTGGGATCTTGAGG |
| qPCR *LjKAI2a* | Sc282 | CGGTGCAGGAGTTTAGCAGA |
| | Sc283 | TCCACAATAGACACGCCACC |
| qPCR *LjKAI2b* | Sc284 | AAGAAAGACCTGGCGGTTCC |
| | Sc285 | CCTCCGTTGACATGACCTCC |
| qPCR *LjDLK2* | MG027 | CTCCTTGGTGCTTCTCCCAG |
| | MG028 | AAAGCCGAAGCCAGTTTTCA |

| | | |
|---|---|---|
| qPCR *LjD14* | D14_qPCR_F | ACAGCGTCCGAGAAAACTC |
| | D14_qPCR_R | AGCAATGGAGGCCAACTAC |

### Supplemental Table 4 | Plasmids.

| **Name** | **Description** |
|---|---|
| **Golden Gate Level 0** | |
| L0 c*LjKAI2a*$_{M160, L190,W157}$ A | PCR amplification of *L. japonicus* Gifu cDNA with primers Sc505 +ST161. Assembly by StuI cut ligation into L0-Amp (BB01) |
| L0 c*LjKAI2a*$_{M160, L190,W157}$ B | PCR amplification of *L. japonicus* Gifu cDNA with primers ST163 +ST164. Assembly by StuI cut ligation into L0-Amp (BB01) |
| L0 c*LjKAI2a*$_{M160, L190,W157}$ C | PCR amplification of *L. japonicus* Gifu cDNA with primers ST165 +Sc498. Assembly by StuI cut ligation into L0-Amp (BB01) |
| L0 c*LjKAI2b*$_{L161,S191,F158}$ A | PCR amplification of *L. japonicus* Gifu cDNA with primers Sc506 +ST169. Assembly by StuI cut ligation into L0-Amp (BB01) |
| L0 c*LjKAI2b*$_{L161,S191,F158}$ B | PCR amplification of *L. japonicus* Gifu cDNA with primers ST171 +ST172. Assembly by StuI cut ligation into L0-Amp (BB01) |
| L0 c*LjKAI2b*$_{L161,S191,F158}$ C | PCR amplification of *L. japonicus* Gifu cDNA with primers ST173 +Sc503. Assembly by StuI cut ligation into L0-Amp (BB01) |
| **Golden Gate Level I** | |
| LI Esp3I p*AtKAI2* A | PCR amplification of *L. japonicus* Gifu genomic DNA with primers Sc232 + Sc233. Assembly by StuI cut ligation into LI-pUC57 (BB02) |
| LI Esp3I p*AtKAI2 B* | PCR amplification of *L. japonicus* Gifu genomic DNA with primers Sc234 + Sc235. Assembly by StuI cut ligation into LI-pUC57 (BB02) |
| LI Esp3I p*AtD14* | PCR amplification of *L. japonicus* Gifu genomic DNA with primers Sc224 + Sc225. Assembly by StuI cut ligation into LI-pUC57 (BB02) |
| LI Esp3I *gAtKAI2* | PCR amplification of *L. japonicus* Gifu genomic DNA with primers Sc238 + Sc239. Assembly by StuI cut ligation into LI-pUC57 (BB02) |
| LI Esp3I *gAtD14* | PCR amplification of *L. japonicus* Gifu genomic DNA with primers Sc237 + Sc238. Assembly by StuI cut ligation into LI-pUC57 (BB02) |
| LI Esp3I *gLjKAI2a* | PCR amplification of *L. japonicus* Gifu genomic DNA with primers Sc243 + Sc244. Assembly by StuI cut ligation into LI-pUC57 (BB02) |
| LI Esp3I *gLjKAI2b* | PCR amplification of *L. japonicus* Gifu genomic DNA with primers Sc246 + Sc247. Assembly by StuI cut ligation into LI-pUC57 (BB02) |
| LI Esp3I *gLjD14* | PCR amplification of *L. japonicus* Gifu genomic DNA with primers Sc240 + Sc241. Assembly by StuI cut ligation into LI-pUC57 (BB02) |
| LI Esp3I *gLjkai2a-1* | PCR amplification of *L. japonicus kai2a-1* genomic DNA with primers Sc243 + ST97 and ST96 +Sc244. Assembly by BpiI and StuI cut ligation into LI-pUC57 (BB02) |
| LI Esp3I *cLjkai2a-1* | PCR amplification of *L. japonicus kai2a-1* coding DNA with primers Sc243 + Sc244. Assembly by StuI cut ligation into LI-pUC57 (BB02) |
| LI Esp3I *cLjKAI2a* | PCR amplification of *L. japonicus* Gifu coding DNA with primers Sc243 + Sc244. Assembly by StuI cut ligation into LI-pUC57 (BB02) |
| LI Esp3I c*LjKAI2b* | PCR amplification of *L. japonicus* Gifu cDNA with primers Sc246 + Sc248. Assembly by StuI cut ligation into LI-pUC57 (BB02) |

| | |
|---|---|
| LI Esp3I c*LjKAI2a*$_{M160,\ L190,W157}$ | Assembled by BpiI cut ligation from: L0 c*LjKAI2a*$_{M160,\ L190,W157}$ A + L0 c*LjKAI2a*$_{M160,\ L190,W157}$ B + L0 c*LjKAI2a*$_{M160,\ L190,W157}$ C + LI-BpiI (BB03) |
| LI Esp3I c*LjKAI2b* $_{L161,S191,F158}$ | Assembled by BpiI cut ligation from: L0 c*LjKAI2b* $_{L161,S191,F158}$ A + L0 c*LjKAI2b* $_{L161,S191,F158}$ B + L0 c*LjKAI2b* $_{L161,S191,F158}$ C + LI-BpiI (BB03) |
| **Golden Gate Level II** | |
| LIIc F 1-2 POI:GOI:*HygroR* | Assembled by BsaI cut ligation from: LI A-B POI (G082) + LI B-C dy (BB06) + LI C-D GOI + LI D-E dy (BB08) + LI E-F nos-T (G006) + LI F-G HygroR (G095) + LIIc F 1-2 (BB30) |
| LIIc R 3-4 p*35S:mCherry* | Assembled by BsaI cut ligation from: LI A-B p*35S* (G005) + LI B-C dy (BB06) + LI C-D *mCherry* (G023) + LI D-E dy (BB08) + LI E-F 35S-T (G059) + LI F-G dy (BB09) + LIIc R 3-4 (BB34) |
| **Golden Gate Level III** | |
| LIIIβ POI:GOI:*HygroR* | Assembled by BpiI cut ligation from: LIIc F 1-2 POI:GOI:HygroR + LII 2-3 ins (BB43) + LIIc R 3-4 p35S*:mCherry* + LII 4-6 dy (BB41) + LIIIβ F A-B (BB53) |
| LIIIβ p*AtKAI2:gAtKAI2* | Assembled by Esp3I cut ligation from: LIIIβ F A-B POI:GOI:HygroR + LI Esp3I *pAtKAI2* A + LI Esp3I *pAtKAI2* A + LI Esp3I *gAtKAI2* |
| LIIIβ p*AtKAI2:gAtD14* | Assembled by Esp3I cut ligation from: LIIIβ F A-B POI:GOI:HygroR + LI Esp3I *pAtKAI2* A + LI Esp3I *pAtKAI2* A + LI Esp3I *gAtD14* |
| LIIIβ p*AtKAI2:gLjKAI2a* | Assembled by Esp3I cut ligation from: LIIIβ F A-B POI:GOI:HygroR + LI Esp3I *pAtKAI2* A + LI Esp3I *pAtKAI2* A + LI Esp3I *gLjKAI2a* |
| LIIIβ p*AtKAI2:gLjKAI2b* | Assembled by Esp3I cut ligation from: LIIIβ F A-B POI:GOI:HygroR + LI Esp3I *pAtKAI2* A + LI Esp3I *pAtKAI2* A + LI Esp3I *gLjKAI2b* |
| LIIIβ p*AtKAI2: gLjkai2a-1* | Assembled by Esp3I cut ligation from: LIIIβ F A-B POI:GOI:HygroR + LI Esp3I *pAtKAI2* A + LI Esp3I *pAtKAI2* A + LI Esp3I *gLjkai2a-1* |
| LIIIβ p*AtKAI2: cLjkai2a-1* | Assembled by Esp3I cut ligation from: LIIIβ F A-B POI:GOI:HygroR + LI Esp3I *pAtKAI2* A + LI Esp3I *pAtKAI2* A + LI Esp3I *cLjkai2a-1* |
| LIIIβ p*AtKAI2:gLjD14* | Assembled by Esp3I cut ligation from: LIIIβ F A-B POI:GOI:HygroR + LI Esp3I *pAtKAI2* A + LI Esp3I *pAtKAI2* A + LI Esp3I *gLjD14* |
| LIIIβ p*AtD14:gAtD14* | Assembled by Esp3I cut ligation from: LIIIβ F A-B POI:GOI:HygroR + LI Esp3I *pAtKD14* + LI Esp3I *gAtD14* |
| LIIIβ p*AtD14:gAtKAI2* | Assembled by Esp3I cut ligation from: LIIIβ F A-B POI:GOI:HygroR + LI Esp3I *pAtKD14* + LI Esp3I *gAtKAI2* |
| LIIIβ p*AtD14:gLjD14* | Assembled by Esp3I cut ligation from: LIIIβ F A-B POI:GOI:HygroR + LI Esp3I *pAtKD14* + LI Esp3I *gLjD14* |
| LIIIβ p*AtD14:gLjKAI2a* | Assembled by Esp3I cut ligation from: LIIIβ F A-B POI:GOI:HygroR + LI Esp3I *pAtKD14* + LI Esp3I *gLjKAI2a* |
| LIIIβ p*AtD14:gLjKAI2b* | Assembled by Esp3I cut ligation from: LIIIβ F A-B POI:GOI:HygroR + LI Esp3I *pAtKD14* + LI Esp3I *gLjKAI2b* |
| **Protein induction** | |
| pSUMO c*LjKAI2a* | PCR amplification from LI Esp3I *cLjKAI2a* with primers MW1002 + MW1003. Assembly by Gibson cloning |
| pSUMO c*LjKAI2b* | PCR amplification from LI Esp3I *cLjKAI2b* with primers MW1002 + MW1004. Assembly by Gibson cloning |
| pSUMO c*LjKAI2a*$_{M160,\ L190,W157}$ | PCR amplification from LI Esp3I *cLjKAI2a (3b)* with primers MW1002 + MW1003. Assembly by Gibson cloning |
| pSUMO c*LjKAI2b* $_{L161,S191,F158}$ | PCR amplification from LI Esp3I *cLjKAI2b (3a)* with primers MW1002 + MW1004. Assembly by Gibson cloning |

| | |
|---|---|
| pSUMO c*LjKAI2a*$_{M160, L190}$ | Rolling circle PCR amplification from pSUMO *LjKAI2a (3b)* with primers Sc604 + Sc605. |
| pSUMO c*LjKAI2b* $_{L161,S191,F158}$ | Rolling circle PCR amplification from pSUMO *LjKAI2b (3a)* with primers Sc606 + Sc607. |
| pSUMO c*LjKAI2*$_{,W157}$ | Rolling circle PCR amplification from pSUMO *LjKAI2a* with primers ST177 + ST178. |
| pSUMO c*LjKAI2b*$_{,F158}$ | Rolling circle PCR amplification from pSUMO *LjKAI2b* with primers ST179 + ST180. |

**S5 Table 5.** Statistical results of ANOVA for multiple comparisons.

| Figure | genotype/treatment/gene | *post hoc* test | p-value | F-value |
|---|---|---|---|---|
| Fig. 2A | - | Tukey | ≤ 0.001 | $F_{14/1438}$ = 125.3 |
| Fig. 2C | - | Tukey | ≤ 0.001 | $F_{11/132}$ = 45.6 |
| Fig. 3B | WT (Ler) | Tukey | ≤ 0.001 | $F_{2/311}$ = 244 |
| | *kai2-2* | | = 0.18 | $F_{2/300}$ = 1.71 |
| | *AtKAI2 #1* | | ≤ 0.001 | $F_{2/122}$ = 31.9 |
| | *AtKAI2 #3* | | ≤ 0.001 | $F_{2/303}$ = 116.4 |
| | *LjKAI2a #10b* | | ≤ 0.001 | $F_{2/316}$ = 65.7 |
| | *LjKAI2a #11b* | | ≤ 0.001 | $F_{2/313}$ = 42 |
| | *LjKAI2b #1b* | | ≤ 0.001 | $F_{2/296}$ = 33.4 |
| | *LjKAI2b #5b* | | ≤ 0.001 | $F_{2/288}$ = 87.4 |
| Fig. 3C | WT (Col) | Tukey | ≤ 0.001 | $F_{2/311}$ = 158.3 |
| | K02821 | | ≤ 0.001 | $F_{2/353}$ = 100.3 |
| | WT (Ler) | | ≤ 0.001 | $F_{2/384}$ = 499.6 |
| | *htl-2* | | ≤ 0.05 | $F_{2/391}$ = 3.2 |
| | #18 | | ≤ 0.001 | $F_{2/383}$ = 104.8 |
| | #23 | | ≤ 0.001 | $F_{2/253}$ = 127 |
| Fig. 3D | WT (Col) | Tukey | ≤ 0.001 | $F_{2/415}$ = 1008 |
| | *d14-1 kai2-2* | | = 0.22 | $F_{2/353}$ = 1.54 |
| | *LjKAI2a #32* | | ≤ 0.001 | $F_{2/287}$ = 50 |
| | *LjKAI2a #46* | | ≤ 0.001 | $F_{2/184}$ = 85 |
| | *LjKAI2b #29* | | ≤ 0.001 | $F_{2/283}$ = 9.4 |
| | *LjKAI2b #31* | | ≤ 0.05 | $F_{2/244}$ = 3.9 |
| Fig. 4B | LjKAI2a | Dunnett | ≤ 0.0001 | $F_{5/12}$ = 96.1 |
| | LjKAI2a$_{M160,L190}$ | | ≤ 0.001 | $F_{5/12}$ = 9.5 |
| | LjKAI2a$_{M160,L190, W157}$ | | = 0.227 | $F_{5/12}$ = 1.63 |
| | LjKAI2a $_{W157}$ | | ≤ 0.05 | $F_{5/12}$ = 4.17 |
| | LjKAI2b | | = 0.632 | $F_{5/12}$ = 0.70 |
| | LjKAI2b$_{L161,M191}$ | | = 0.001 | $F_{5/12}$ = 8.9 |
| | LjKAI2b $_{L161,M191,F158}$ | | ≤ 0.0001 | $F_{5/12}$ = 56.9 |
| | LjKAI2b$_{F158}$ | | ≤ 0.0001 | $F_{5/12}$ = 29.54 |
| Fig. 6C | - | Tukey | ≤ 0.001 | $F_{6/103}$ = 35 |
| Fig. 6D | - | Tukey | ≤ 0.001 | $F_{4/67}$ = 19.9 |

| | | | | |
|---|---|---|---|---|
| Fig. 6E | - | Tukey | ≤ 0.001 | $F_{6/605} = 26.5$ |
| Fig. 7A | KAR1 PRL | Tukey | ≤ 0.001 | $F_{3/209} = 7.40$ |
| | KAR1 PER | | ≤ 0.001 | $F_{3/209} = 11.1$ |
| | KAR1 PER density | | ≤ 0.01 | $F_{3/209} = 5.51$ |
| | KAR2 PRL | | = 0.51 | $F_{3/217} = 0.77$ |
| | KAR2 PER | | = 0.18 | $F_{3/217} = 1.64$ |
| | KAR2 PER density | | = 0.72 | $F_{3/217} = 0.44$ |
| | *rac*-GR24 PRL | | = 0.74 | $F_{3/203} = 0.42$ |
| | *rac*-GR24 PER | | = 0.07 | $F_{3/203} = 2.45$ |
| | *rac*-GR24 PER density | | = 0.43 | $F_{3/203} = 0.92$ |
| Fig. 7B | - | Dunnett | ≤ 0.01 | $F_{3/188} = 4.08$ |
| Fig. 7C | WT | Tukey | ≤ 0.001 | $F_{2/9} = 30.7$ |
| | *max2-4* | | = 0.20 | $F_{2/9} = 1.97$ |
| Fig. S8A | *KAI2a* | Tukey | ≤ 0.001 | $F_{5/18} = 39.5$ |
| | *KAI2b* | | ≤ 0.001 | $F_{5/18} = 33.7$ |
| Fig. S9D | - | Tukey | ≤ 0.001 | $F_{9/714} = 178.8$ |
| Fig. S10B | KAR1 | Tukey | ≤ 0.001 | $F_{3/396} = 33.1$ |
| | KAR2 | | ≤ 0.001 | $F_{3/390} = 16.5$ |
| | *rac*-Gr24 | | ≤ 0.001 | $F_{3/392} = 35$ |
| Fig. S10C | WT | Dunnett | ≤ 0.001 | $F_{2/313} = 30$ |
| | *kai2a-1* | | = 0.08 | $F_{2/234} = 2.51$ |
| | *kai2b-1* | | ≤ 0.001 | $F_{2/302} = 29.3$ |
| | *kai2b-3* | | ≤ 0.001 | $F_{2/308} = 14.2$ |
| | *kai2a-1 kai2b-1* | | = 0.99 | $F_{2/272} = 0.01$ |
| Fig. S10D | WT | Dunnett | ≤ 0.001 | $F_{2/246} = 51$ |
| | *max2-4* | | = 0.25 | $F_{2/204} = 1.38$ |
| Fig. S10E | WT | Dunnett | ≤ 0.001 | $F_{3/8} = 28.4$ |
| | *kai2a-1* | | ≤ 0.001 | $F_{3/8} = 53$ |
| | *kai2b-3* | | ≤ 0.001 | $F_{3/8} = 26$ |
| | *kai2a-1 kai2b-1* | | ≤ 0.001 | $F_{3/8} = 105.8$ |
| | *max2-4* | | = 0.99 | $F_{3/8} = 0.04$ |
